# Systemic Infection Facilitates Transmission of *Pseudomonas aeruginosa*

**DOI:** 10.1101/765339

**Authors:** Kelly E. R. Bachta, Jonathan P. Allen, Bettina H. Cheung, Cheng-Hsun Chiu, Alan R. Hauser

## Abstract

Health care-associated infections such as *Pseudomonas aeruginosa* (PA) bacteremia pose a major clinical risk for hospitalized patients, and efforts to limit them are a priority. The fitness pressures accounting for PA virulence factors that facilitate bloodstream infections are unclear, as these infections are presumed to be a “dead-end” and have no impact on transmission. Here, we used a mouse model to show that PA spreads from the bloodstream to the gallbladder, where it replicates to extremely high numbers. Bacteria in the gallbladder then seed the intestines and feces, leading to transmission to uninfected cage-mate mice. The findings demonstrate that the gallbladder is critical for spread of PA from the bloodstream to the feces during bacteremia, a process that promotes transmission.

## MAIN

Annually in the United States, 440,000 adults contract a health care-associated infection (HAI) resulting in increased mortality for individual patients and an estimated $10 billion in health care costs^1^. While progress has been made in minimizing patient-to-patient spread of HAI bacteria, much remains unknown regarding how these pathogens are transmitted between patients. *Pseudomonas aeruginosa* (PA) is a common cause of HAI, and in a head-to-head comparison of bloodstream infections (bacteremias), PA was associated with higher mortality than other bacteria^2^. This, coupled with the rapid emergence of multidrug-resistant (MDR) and extensively drug-resistant (XDR) PA, make prevention of PA transmission critical^3,4^. However, such prevention has been hampered by an incomplete understanding of in-host PA infection dynamics and their relationship to transmission among hospitalized patients.

Clinically, PA bloodstream infections are regarded as “dead-ends”, meaning that the bacteria in the blood are neither transmitted to other patients or spread back to the environment. Once bacteria reach the bloodstream, they are either cleared by the host with the help of antibiotics or die with the host if the infection is fatal. In either case, genetic adaptations in PA that confer an enhanced ability to disseminate to the bloodstream are subsequently eliminated from the gene pool. In this regard, PA is thought to differ from gastrointestinal pathogens such as *Salmonella typhi*^5–7^ and *Listeria monocytogenes*^8–10^, which have evolved complex lifecycles that include trafficking from the blood to the gallbladder and subsequently to the intestines, from which they are excreted in feces to facilitate transmission to new hosts. In contrast, PA bloodstream infections would appear to be accidents from which the bacteria gain no benefit. Thus, the fitness pressures responsible for the evolution of virulence factors that facilitate dissemination of PA to the bloodstream are poorly understood.

To address these issues, we examined the fate of PA bacteria in the bloodstream using a murine model in conjunction with sequence tag-based analysis of microbial populations (STAMP)^9,11^. STAMP utilizes barcoded wild-type strains in combination with deep sequencing to track populations of bacteria, assess infection bottlenecks, and measure the relatedness of pathogen populations at different sampling sites. We demonstrated that bloodstream PA bacteria were trapped by the liver and spread to the gallbladder, where they unexpectedly replicated to extremely high numbers. The bacterial population in the gallbladder seeded the intestines and was excreted in the feces in a manner that facilitated spread to cage-mate mice. Mice lacking a gallbladder (cholecystectomized) excreted dramatically fewer PA bacteria in their intestinal tracks than wild-type mice, highlighting a critical role for this organ in excretion and environmental contamination. The liver-gallbladder-intestinal excretion pathway may therefore facilitate spread of PA from the bloodstream to the intestines and simultaneously allow amplification of PA. In this way, PA bacteria from systemic infections are returned to the environment in high numbers, where they may facilitate transmission.

### PA disseminates to the gallbladder and gastrointestinal tract following systemic infection

We examined the fate of PA during a bloodstream infection utilizing a mouse model of bacteremia in which mice naïve to antibiotic treatment (wild-type) were infected via tail vein to allow for consistent and reproducible delivery of bacteria directly into the bloodstream. To define the pattern of dissemination following PA bloodstream infection, we engineered a PA clinical isolate (PABL012) for *in vivo* bioluminescence imaging using this model. Mice infected with PABL012*_lux_* by tail vein injection displayed an intense bioluminescent focus in their ventral mid-section by 24 hours (Fig. 1A, B), which was localized to the gallbladder (Fig. 1C, G). Moderate bioluminescence signals were observed in the stomach (Fig. 1H) and liver (Fig. 1F), while weak signals were detected in the lungs (Fig. 1D), spleen (Fig. 1E) and intestinal tract (Fig. 1I).

**Figure 1.**
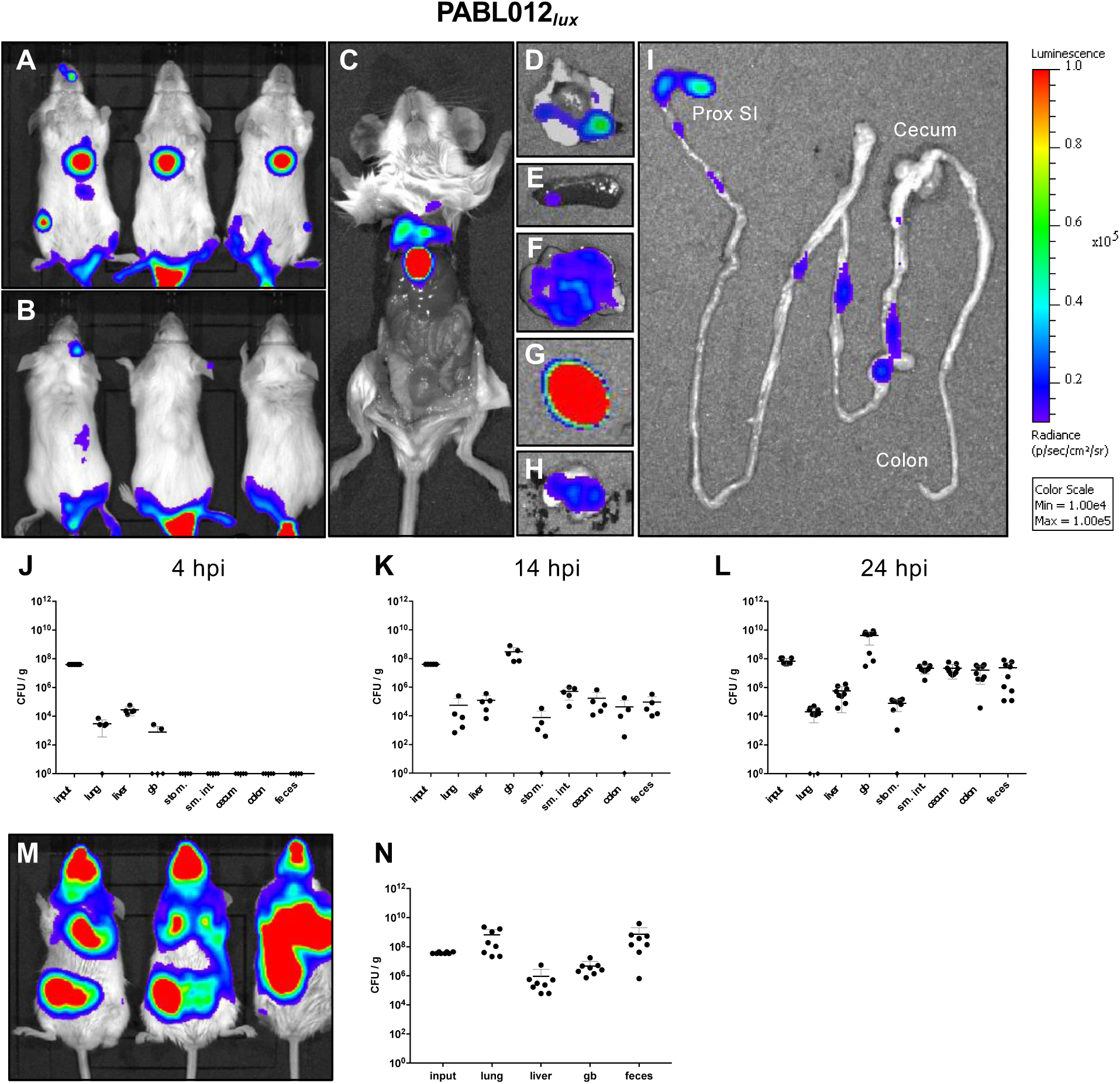
*P. aeruginosa* disseminates to the gallbladder and intestines following bloodstream infection and pneumonia. Mice (n=3) were intravenously injected with ∼2 x 106 CFU of a luciferase expressing strain of *P. aeruginosa*, PABL012*lux*. Ventral (A) and dorsal (B) views of live mice at 24 hours post-infection (hpi) were obtained using IVIS. Mice were subsequently euthanized, the abdominal wall dissected away and internal organs visualized (C). The heart and lung (D), spleen (E), liver (F), gallbladder (G), stomach (H) and remaining intestinal tract (I) were separately removed and imaged to visualize bacterial localization. (Since bioluminescence is dependent on oxygen, the signal present in the anaerobic portions of the intestinal tract may underestimate bacterial numbers.) The scale for all images represents radiance (photons/sec/cm^2^/sr) with minimum and maximum values normalized to 1 x 10^4^ and 1 x 10^5^ respectively. Intravenously infected mice housed on standard bedding were euthanized at 4 hpi (J, n=4), 14 hpi (K, n=5), or 24 hpi (L, n=9). Bacteria were enumerated from several anatomical sites by plating serial dilutions of organ homogenates. (M) BALB/c mice (n=8) were infected with PABL012*_lux_* by intranasal inoculation of ∼2 x 10^6^ CFU and imaged at 24 hpi using IVIS. (N) Mice were subsequently euthanized and bacteria were enumerated from several anatomical sites. Density (CFU per gram [CFU/g] of organ weight) of recovered bacteria are presented (black circles). Organs where no bacteria were recovered are denoted with a diamond on the x-axis. Geometric means (horizontal lines) and SD (whiskers) are shown. (gb) gallbladder, (stom.) stomach, (sm. int.) small intestine.

Given that quantifying bacterial population sizes from *in vivo* bioluminescence is imprecise, additional studies were performed to directly enumerate bacterial counts from infected organs. At 4 hours post-infection (hpi), PA was detected in the liver, lungs, and some of the gallbladders, but no bacteria were detected in the intestinal tract (Fig. 1J). This was followed by a drastic six-logarithm expansion of bacteria in the gallbladder by 14 hpi and similar four- to six-logarithm expansion in the intestinal tract and feces (Fig. 1K). In contrast, total bacterial counts in the lung and liver rose by only one- to two-logarithms over the same period, demonstrating that rapid increases in PA numbers were not a universal phenomenon. Bacterial recovery in the gallbladder reached higher numbers than were originally in the inoculum, suggesting that the gallbladder may be a hospitable niche for PA replication. In support of this, we also observed that PA growth in *ex-vivo* bile preparations mirrored that of enriched medium (Extended Data Fig. 1). By 24 hpi, this dramatic expansion of bacterial populations had plateaued with only slight additional increases in all organs (Fig. 1L). Lumenal contents of the intestinal tract contained similar bacterial CFU as observed in whole intestine homogenates (Extended Data Fig. 2G; compare to Fig. 1L), indicating that most PA bacteria were in the lumen rather than in the bowel wall. These findings indicate that bacteremia is followed by a dramatic expansion of the bacterial population in the gallbladder and intestines, which is accompanied by PA excretion in the feces.

To determine whether fecal shedding was a characteristic of PA, we tested an additional panel of non-clonal isolates including the commonly used laboratory strains PAO1 and PA14, the MDR clinical isolate PABL046, as well as 11 additional clinical isolates (Table 1). All PA strains shared the same pattern of dissemination to the gallbladder and excretion into the feces (Extended Data Figs. 2A-D, 3) despite being phylogenetically distinct, globally distributed, and representing 9 different multilocus sequence types (Extended Data Fig. 3, Table 1). We also challenged male BALB/c mice (Extended Data Fig. 2E) to explore possible gender bias and female C57/BL6 mice (Extended Data Fig. 2F) as an alternate mouse strain background. Male BALB/c mice and female C57/BL6 mice both revealed similar patterns of dissemination, gallbladder expansion, and fecal excretion to the female BALB/c mice. These findings suggest that trafficking to and expansion in the gallbladder, transit to the intestines, and excretion in feces may be universal features of PA bacteremia regardless of PA isolate, murine gender or genetic background.

### The PA population shed in the feces originates in the gallbladder

The above data indicated that significant numbers of PA accumulated in the gallbladder, but questions remained as to how systemic PA gained access to the gallbladder and whether they indeed replicated to high numbers there. To investigate these questions, we utilized an approach to genetically track bacterial populations referred to as sequence tag-based analysis of microbial populations (STAMP)^11^. We generated a library of barcoded but otherwise isogenic PABL012 bacteria (PABL012*_pool_*) containing ∼4000 unique short (∼30 bp) sequence tags inserted into a neutral site on the chromosome. By measuring and comparing barcode frequencies from bacterial populations in different anatomical locations, we could mathematically estimate the founding population size (*N_b_*) at a given sampling site. The *N_b_* value estimates population diversity and reflects how host barriers may shape bacterial populations during systemic infection. Control experiments demonstrated that barcodes did not influence growth rate (Extended Data Fig. 4A) and were stable in the absence of antibiotic selection (Extended Data Fig. 4B). Next, we artificially simulated bottlenecks under controlled *in vitro* conditions by sampling serial 10-fold dilutions of the PABL012*_pool_*. The sizes of these samples were empirically measured by bacterial enumeration (CFU) and in parallel calculated using the STAMP approach and the barcode frequencies in PABL012*_pool_* (*N_b_*) (Extended Data Fig. 4C). The calculated founding population size (*N_b_*) underestimated the actual founding population size, so a calibration curve was generated to adjust our *in vivo* experimental results to account for this underestimation, yielding a value referred to as *N_b_*ʹ (see Methods section for details)^11^.

Mice (n=10) were intravenously injected with PABL012*_pool_*, and per-organ total CFU counts recovered were consistent with those observed in prior experiments (Fig. 2A, C compared to Fig. 1K, L). At 14 and 24 hpi, the mean founding population sizes (*N_b_*ʹ) were quite low (between 35-70 at 14 hpi and 40-60 at 24 hpi) in the majority of the organs. In contrast, the spleen had a larger mean *N_b_*ʹ of 1509 at 14 hpi (Fig. 2A) and 523 at 24 hpi (Fig. 2C), indicative of a more diverse founding population that likely reflects the role of the spleen in filtering bacteria out of the bloodstream^12^. The low *N_b_*ʹ values of the gallbladder and intestines suggest that PA passes through a severe bottleneck in the process of reaching these organs. This low founding population (*N_b_*ʹ) coupled with high bacterial numbers (CFU), suggests that the few PA cells that traverse this narrow bottleneck subsequently replicate to high numbers upon arrival in the gallbladder.

**Figure 2.**
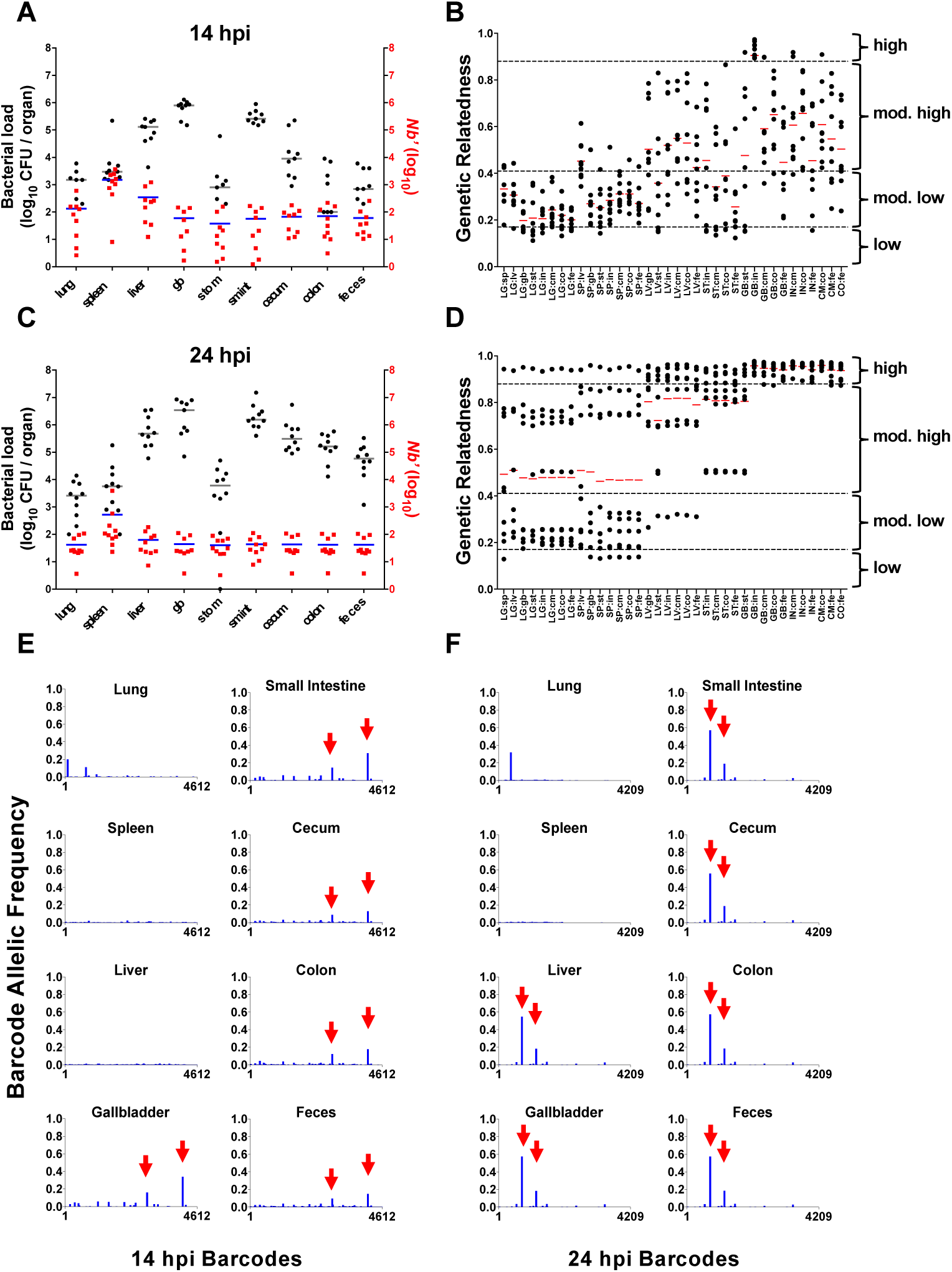
Diversity and relatedness of disseminated *P. aeruginosa* populations following bloodstream infection. BALB/c mice were intravenously injected with ∼2 x 10^6^ CFU of PABL012*_pool_*. (A) Bacterial loads (total CFU, black circles) and *N_b_*′ sizes (red squares) in different tissues were determined at 14 hpi (A) and 24 hpi (C). CFU values are expressed per organ for the lung, spleen, liver, gallbladder (gb), stomach (stom), small intestine (sm int), cecum, colon, and feces. Each black circle and red square represent an organ from one mouse (14 hpi n=9, 24 hpi n=10). Geometric means (horizontal lines) are shown. Genetic relatedness of PA populations at 14 (B) and 24 hpi (D). Each circle represents one organ in pairwise comparison with another organ from the same mouse. Geometric means (red, horizontal bars) are shown. Dashed lines denote genetic relatedness categories which are defined as high (≥0.88), moderately high (<0.88, ≥0.41), moderately low (<0.41, ≥0.17), and low (<0.17). LG (lung), SP (spleen), LV (liver), ST (stomach), GB (gallbladder), IN (small intestine), CM (cecum), CO (colon), FE (feces). GR is impacted in large part by the low allelic frequency barcodes. The relative allelic frequencies of the all unique barcodes in every organ of a representative mouse is shown at (E) 14 hpi (n=4612) and (F) 24 hpi (n=4208). Red arrows denote the top 2 dominant tags in the LV, GB, IN, CM, CO, and FE.

Interestingly, the founding population sizes for PA isolated from the liver, gallbladder, intestines and feces were all quite low, suggesting that these bacterial populations may be related to one another. We hypothesized that the liver may be imposing the tight bottleneck observed in our *N_b_*ʹ calculations. Because of the anatomic link between the liver, gallbladder and intestines, those PA cells that escape the liver and make it to the gallbladder would then undergo unimpeded replication and escape from the host through the intestinal tract. To address this hypothesis, specific barcode frequencies (allelic frequency) from all organs within a given mouse were compared to determine the genetic relatedness (GR) of bacterial populations between two organs in that mouse according to the method of Cavalli and Sforza^13^ (Fig. 2B, D). Basically, this approach relies on the premise that if bacterial populations from two different organs share an origin, the same set of barcodes will be over-represented in each. The PA populations within the gallbladder, small intestine, cecum, colon, and feces of each mouse increased from moderately highly related at 14 hpi (Fig. 2B) to virtually identical at 24 hpi (GR values approaching 1, Fig. 2D). At both 14 and 24 hpi, these populations were dominated by the same relatively few sequence barcodes (Fig. 2E, F, Extended Fig. 5A, 6A), indicating that the same bacterial population had spread from the gallbladder to the intestinal tract and the feces. Interestingly, the populations in the stomach at 14 hpi were poorly related (mod. low) to all other organs suggesting that mice were ingesting fecal pellets from other mice within the same cage (thus ingesting different barcodes). However, at 24 hpi, the populations in the stomach were most closely related to the populations in the liver, gallbladder and intestine which could reflect both ongoing ingestion of fecal pellets as well as reflux of barcodes from those expelled into the small intestine by the gallbladder. Taken together, these data suggest that during bacteremia a small subpopulation of PA accessed the liver and escaped to the gallbladder. Once in the gallbladder this population then underwent a profound expansion, disseminated to the intestines and was excreted in the feces. The lack of high levels of relatedness between the populations in the lung, spleen and liver suggests that these organs were seeded independently by unique founding populations and that the dynamics of population control within each organ were different (Fig. 2B, D).

### PA is shed in the feces of bacteremic mice for up to 10 days

We next examined the duration of PA excretion following bloodstream infection. Mice were infected with a subclinical infectious dose (∼8 x 10^5^ CFU) of PABL012*_lux_*. By 2 days post-infection, 100% of infected mice shed between 10^4^ and 10^8^ CFU of PABL012*_lux_* per gram of feces, and excretion was sustained over 5 days (Fig. 3A). Beginning at day 6 and continuing through the remainder of the experiment, the number of infected mice shedding this concentration of PABL012*_lux_* dramatically declined until all infected mice cleared PA from their stools by day 10 (Fig. 3A). The levels of PA in the feces of infected mice were not influenced by the coprophagic behavior of mice, as infected mice housed on raised wire floors (whereby fecal pellets are inaccessible for ingestion) showed no difference in the fecal bacterial load compared to infected mice housed on normal bedding (Fig. 3C). To further verify that the observed fecal shedding of bacteremic mice was not a consequence of coprophagia, we tested whether PA delivered by orogastric inoculation of bacteremic mice was detectable in feces. A PA strain marked with a chromosomal gentamicin resistance cassette (PABL012*_GM_*) was administered by oral gavage to mice systemically infected either 4 or 10 hours prior with PABL012*_lux_* (Fig. 3D). The orally administered strain could only be detected in the feces of two mice when administered 10 hours after initial injection of PABL012*_lux_* (Fig. 3D). Furthermore, this recovered population of PABL012*_GM_* was 1,000 times smaller than the fecal population of systemically delivered bacteria (PABL012*_lux_*) and had a minimal contribution to the overall bacterial counts in the feces. Together, these experiments indicate that bacteremic mice can shed substantial amounts of PA into their environment for prolonged periods independent of mouse coprophagic ingestion.

**Figure 3.**
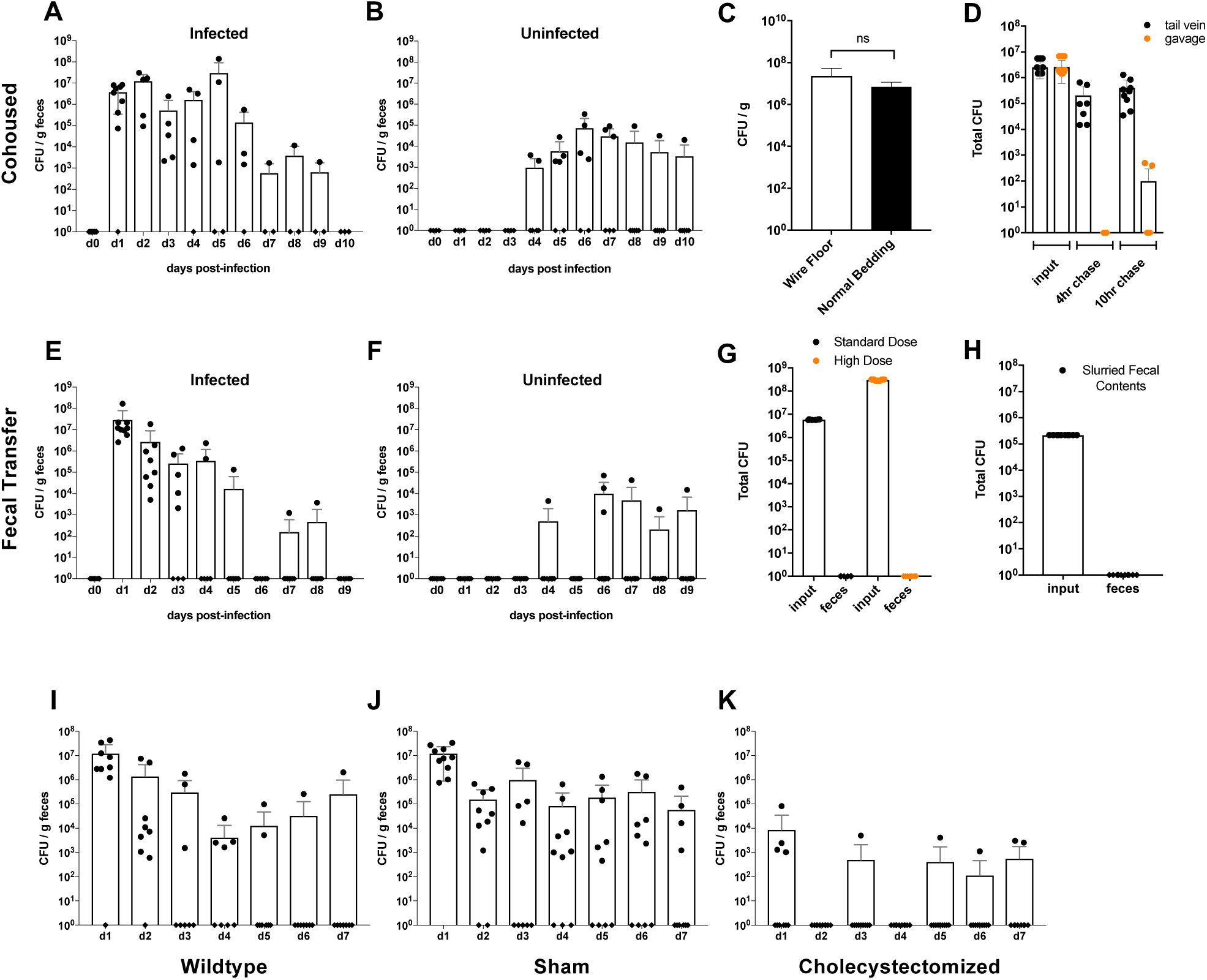
The gallbladder promotes prolonged gastrointestinal shedding of *P. aeruginosa* following bloodstream infection. Three groups of three 6- to 8-week-old BALB/c mice were injected with a sublethal dose (∼8 x10^5^ CFU) of PABL012*_lux_* (A) and co-housed with 2 uninfected cage mates (B) for 10 days (total n=9 infected, n=6 uninfected). Each day, mice were transferred to fresh cages with clean food and water, and fecal pellets were collected from individual mice. Data are presented as the CFU of PA shed per gram feces (CFU/g) where each circle represents data from one mouse. (C) BALB/c mice intravenously infected with ∼2 x 10^6^ CFU of PABL012*_lux_* were housed on normal bedding (n=5) or raised wire racks (n=10), and PA was measured in the feces at 24 hpi. (ns) not significant by Mann-Whitney test. (D) Mice (n=9) were infected intravenously with ∼2 x 10^6^ CFU of PABL012*_lux_* (black circles), followed 4 hours or 10 hours later with oral gavage of ∼2 x 10^6^ CFU of PABL012*_LacZ_* (orange circles) and CFU/g feces reported at 24 hpi. (E) All shed fecal pellets from infected mice (n=8) were transferred daily to the cages of (F) uninfected mice (n=9) and fecal shedding monitored for 9 days. Data are presented as CFU/g feces of PA shed, where each circle represents one mouse. (G) Uninfected BALB/c mice were orally administered standard (∼2 x 10^6^ CFU, n=6, black circles) or high (∼3 x 10^8^ CFU, n=8, orange circles) doses of PABL012*_LacZ_* by gavage, and PA levels were measured in the feces at 24 hpi. (H) Fecal pellets from previously infected mice (n 25) were homogenized, concentrated and orally administered (∼2 x 10^5^ CFU) to uninfected mice by gavage. PA was measured in the feces at 24 hpi. (I, J, K) Two groups of 5- to 7-week-old BALB/c mice (n=5, 2 replicates) underwent cholecystectomy and sham surgery. Mice following sham surgery (J) and mice following cholecystectomy (K) were allowed to recover for one week post-operatively, and they, in addition to age-matched mice that had not undergone surgery (I), were subsequently challenged intravenously with ∼2 x 10^6^ CFU of PABL012*_lux_*. Fecal pellets were collected daily from each group. Data are presented as CFU/g feces. On shedding day 1, P-values were determined for wild type vs. sham (*p* = 0.9, ns), wildtype vs. cholecystectomized (*p* = 0.004, **), and sham vs. cholecystectomized (*p* = 0.0005, ***) mice via ANOVA analysis with Kruskal-Wallis test for multiple corrections. For scatter plots, each circle (or diamond) represents data from one mouse. When no bacterial CFU were recovered, data are represented as diamonds on the x-axis. Geometric means (boxes), and SD (whiskers) are shown.

### Fecal PA is transmitted to uninfected mice

We next examined the consequences of such high levels of PA fecal excretion into the local environment following bloodstream infection. Given that PA infections in BALB/c mice require invasive administration of the bacteria, we hypothesized that uninfected mice would not develop any signs of an active infection when co-housed with systemically infected cage-mates. In addition, the data presented above suggested that the gastrointestinal (GI) tract of uninfected mice was impervious to the passage of PA following orogastric inoculation, so we anticipated that ingestion of PA-laden feces would not enable transmission of PA to cage-mate mice. Mice infected intravenously with PABL012*_lux_* (∼8 x 10^5^ CFU) were co-housed with uninfected mice, and both sets of mice were monitored daily for outward signs of illness and fecal excretion of PA. Daily cage and water changes were performed to minimize any potential environmental PA reservoirs. As hypothesized, no uninfected mice displayed any outward signs of PA infection; however, we observed that approximately 60% of the uninfected mice began to excrete PABL012*_lux_* (10^3^ - 10^4^ CFU/g feces) by day 4, with peak fecal excretion by uninfected mice on day 6 (Fig. 3B). This was followed by a steady decline in bacterial counts in the feces over the remaining 4 days. Moreover, one uninfected mouse continued to shed PABL012*_lux_* at the end of the experiment despite clearance from the originally infected animals suggesting that this mouse maintained prolonged carriage of PA in the GI tract. These experiments revealed that PA shed by systemically infected mice may facilitate transmission to uninfected co-housed mice that is detectable in the feces.

We hypothesized that PABL012*_lux_* detected in the feces of uninfected mice came from ingestion of contaminated fecal pellets shed by infected mice. This observation was somewhat surprising, given our previous results demonstrating the negligible impact of bacterial inoculation by oral gavage (Fig. 3D). Likewise, previous work in mouse models demonstrated that PA cannot access the GI tract through orogastric routes unless extremely high doses of bacteria (>10^7^ CFU/mL drinking water) were administered to antibiotic-treated animals depleted of GI microbiota^14–16^. We further investigated this phenomenon by delivering both standard (6 x 10^6^ CFU) and high doses (3 x 10^8^ CFU) of media-grown PABL012*_lux_* to mice by oral gavage. This type of inoculation resulted in no detectable PA in the feces at 24 hours (Fig. 3G). To test whether the material and microbes present in feces could facilitate PA carriage, we homogenized PABL012*_lux_*-contaminated fecal pellets into a slurry and orogastrically inoculated the slurry (2 x 10^5^ CFU, maximum obtainable dose from pellets of ≈25 shedding mice) into healthy mice. No bacteria could be detected in the feces of these mice at 24 hours post gavage (Fig. 3H), indicating that the presence of contaminated fecal material is not sufficient to allow orally administered PA to result in carriage. We therefore hypothesized that the nature of the fecal pellet in some way facilitates this process. We suspect that the deeper portions of the pellet could enhance the survival of anaerobic bacteria that promote PA carriage in the GI tract or that pellet architecture might enable PA survival during passage through the acidic environment of the stomach. To address this, we collected all fecal pellets shed daily from mice infected intravenously with PABL012*_lux_* (infectious doses not determined but estimated to be significantly less than the orogastric doses described above) and transferred only the contaminated fecal pellets to the cages of uninfected mice simultaneously with daily cage and water changes. One uninfected mouse began to shed PABL012*_lux_* by day 4 and additional mice by day 6 (10^3^ - 10^4^ CFU/g feces, Fig. 3E, F). These findings confirm that exposure to PA contained within fecal pellets is sufficient to facilitate transmission to uninfected animals and suggest that these pellets in some manner facilitate this transmission. In summary, mice systemically infected with PA are capable of contaminating their environment through fecal shedding, which in turn allows transmission to naïve mice that have ingested the fecal pellets.

### The gallbladder is crucial for fecal excretion of PA following bloodstream infection

Given the results of our STAMP analysis indicating that the population of PA excreted in the feces was preceded by dramatic expansion in the gallbladder, we wondered what impact removal of the gallbladder would have on this excretion. Wild-type, sham-surgery (peritoneum opened and repaired), and cholecystectomized mice (n=5, 2 replicates) were infected with the standard dose (2 x 10^6^ CFU) of PABL012*_lux_*. Daily cage and water changes were performed to minimize any potential environmental contamination. On day 1, 90% of the wild-type and 100% of the sham mice excreted high levels of PABL012*_lux_* (between 10^6^ and 10^8^ CFU/g feces), whereas only 40% of the cholecystectomized mice excreted any PABL012*_lux_* and at significantly lower levels (between 10^3^ and 10^5^ CFU/g feces). The proportion of shedding mice in all groups decreased through day 7, but the wild-type and sham mice continued to excrete higher levels of PA than cholecystectomized mice (Fig. 3I, J, K). These results indicate that the bacterial expansion of PA in the gallbladder is crucial for high levels of bacterial excretion into the environment and may play a role in transmission.

### Type III secretion dramatically impacts PA dissemination and injury to the gallbladder

We sought to determine whether the high numbers of PA observed in the gallbladder resulted in damage to the epithelium of this organ and whether known PA virulence factors played a role in this process. To do so, we examined gallbladders of infected animals for histologic and ultrastructural evidence of injury. Gallbladders from mock-infected animals, having no visible bacteria in the lumen, had a typical mucosal layer consisting of columnar epithelial cells with basal nuclei, extended microvilli, and an intact lamina propria (Fig. 4A, B). In contrast, the mucosa of gallbladders from infected mice displayed a general distortion of the cellular architecture characterized by large nuclei, widened tight junctions, increased vacuoles, and disrupted lamina propria (Fig. 4A, B). Disrupted microvillus architecture, characterized by blunted, disordered, or absent microvilli, was observed at the epithelial surface, especially in areas of close association with PA bacteria (Fig. 4B). In mice infected with PABL012*_lux_*, we also noted intermittent focal bacterial aggregates at the epithelial surface overlying damaged columnar epithelial cells (Fig. 4A, *Pa*). Neutrophils were observed at both the basolateral surface (arrows) of the epithelial layer and extravasating through it towards visible lumenal bacteria (arrowhead) (Fig. 4C).

**Figure 4.**
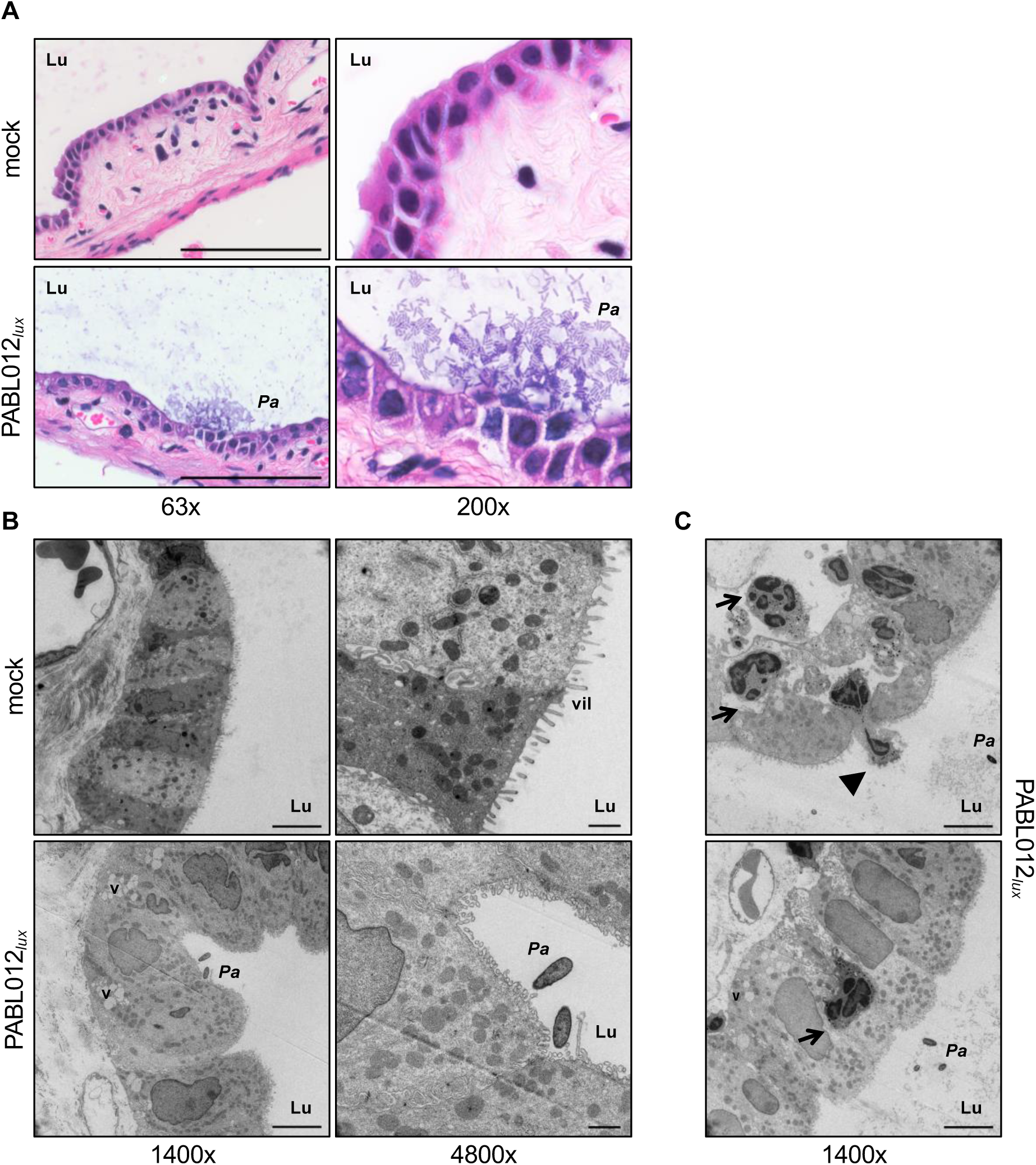
*P. aeruginosa* infection damages the gallbladder epithelium. BALB/c mice intravenously infected with PABL012*_lux_* (∼2 x 10^6^ CFU) were euthanized at 24 hpi to harvest gallbladders for histological preparation. (A) Hematoxylin & eosin (H&E) staining of mock (PBS) or PABL012*_lux_*-infected mouse gallbladder sections at 63x and 100x (scale bar = 100 μM). (B, C) TEM of gallbladder sections from mock (PBS) or PABL012*_lux_*-infected mice (scale bars = 5 μM (1400x), 1 μM (4800x)). Black arrows indicate neutrophils, the black arrowhead shows a transmigrating neutrophil extravasating into the gallbladder lumen. (Lu) lumen, (V) vacuoles, (vil) villi, *P. aeruginosa* (*Pa*).

The type III secretion system (T3SS) is a major driver of virulence in PA and is known to damage epithelial surfaces^17,18^ through direct injection of cytotoxic effector proteins. To assess the role of the T3SS in expansion within the gallbladder and intestines, mice were infected with a T3SS mutant (PABL012Δ*pscJ_lux_*) and monitored for disease progression. The T3SS mutant was absent from the liver, gallbladder, and feces when infected at the standard wild-type dose (Fig. 5A, B). In addition, the PABL012Δ*pscJ_lux_* mutant phenotype could not be rescued by coinfection with wild-type PABL012*_lux_* (Fig. 5C), suggesting that any *in vivo* bottlenecks act directly on the bacteria. The attenuation of the T3SS mutant could be overcome by increasing the infectious dose 100-fold (Fig. 5A, B). However, in contrast to the mucosal damage observed with wild-type infection, no damage to the gallbladder epithelium was observed at this higher dose (Fig. 5D). Thus, an intact T3SS is important for PA dissemination to the gallbladder and subsequent fecal shedding, and results in damage to the gallbladder epithelium.

**Figure 5.**
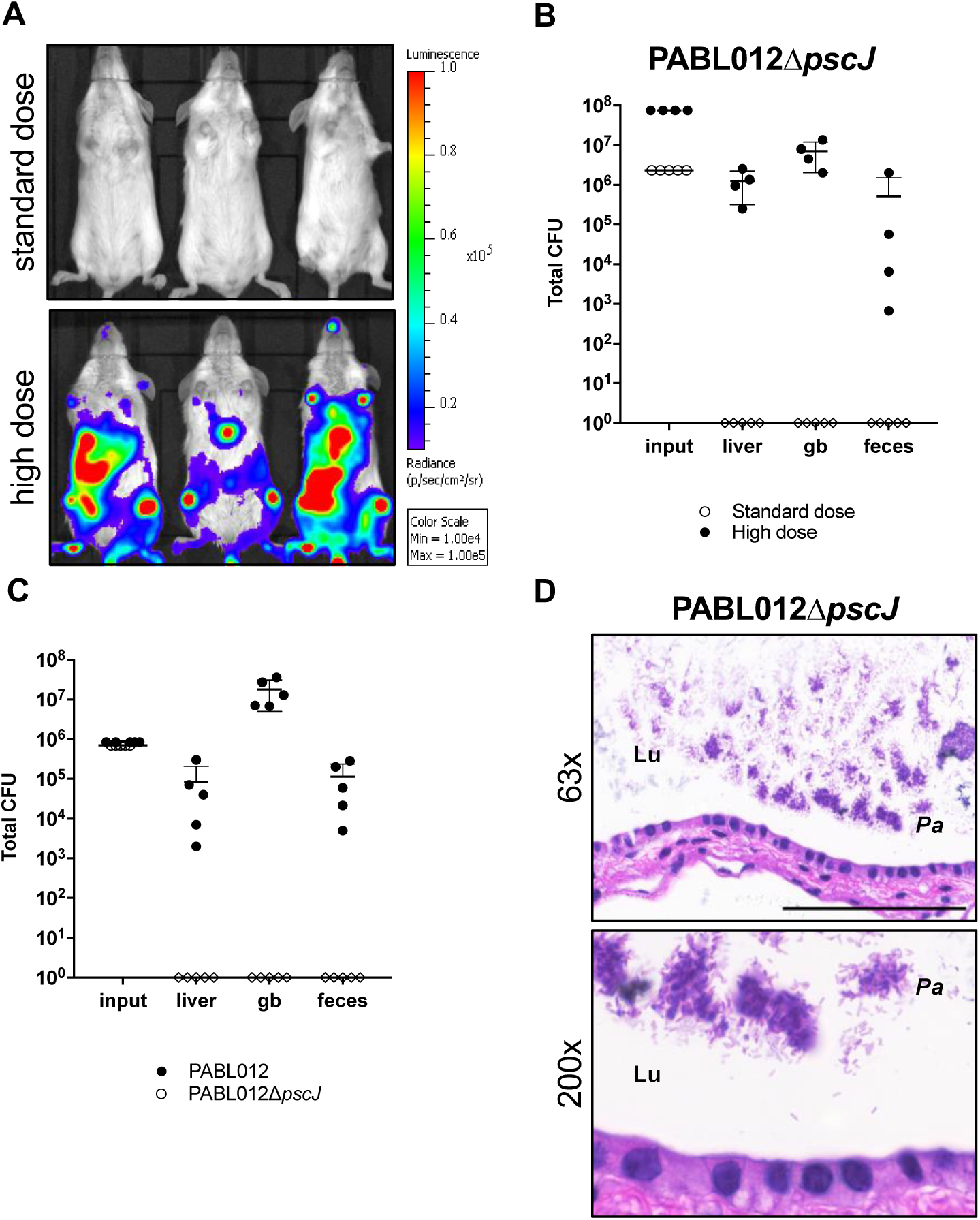
The *P. aeruginosa* type III secretion system (T3SS) contributes to gastrointestinal shedding. (A) BALB/c mice were infected intravenously with a standard (∼2 x 10^6^ CFU, open circles) or a high dose (∼8 x 10^7^ CFU, filled circles) of a PABL012Δ*pscJ_lux_* T3SS mutant and imaged at 24 hpi using IVIS. (B) Mice (n=5) were subsequently euthanized and bacteria enumerated from the liver, gallbladder (gb), and feces. (C) BALB/c mice (n=5) were co-infected intravenously with a 1:1 mix totaling ∼2 x 10^6^ CFU of PABL012*_lacZ_* (filled black circles) and PABL012Δ*pscJ_lux_* (open circles and diamonds). Liver, gb, and feces were harvested at 24 hpi for enumeration of viable bacteria. For scatter plots, each circle or diamond represents data from one mouse. Where no CFU were recovered, data are represented as diamonds on the x-axis. (D) H&E stained sections of gallbladders from mice infected with the high dose of PABL012Δ*pscJ_lux_*. (Lu) lumen, *P. aeruginosa* (*Pa*). Scale bar represents 100 μM (63x).

### PA disseminates to the gallbladder and is shed in the feces following pneumonia

While PA is a frequent cause of bloodstream infections, it is also an important cause of ventilator-associated pneumonia^19^ and previous studies have demonstrated that PA is capable of escaping into the bloodstream during pneumonia^20^. Given our observations following direct inoculation of PA into the bloodstream, we hypothesized that, following pneumonia, PA would disseminate to and expand within the gallbladder. To determine whether gallbladder expansion and fecal shedding were observed following PA pneumonia, 6 to 8-week old BALB/c mice were infected by intranasal inoculation with PABL012*_lux_* (standard dose, 2 x 10^6^ CFU) and monitored for 24 hpi. Infected mice displayed a largely disseminated pattern of infection (Fig. 1M) and contained high numbers of bacteria in the liver, gallbladder and feces (Fig. 1N). These data indicate that following pneumonia PA also expands within the liver and gallbladder, and is ultimately shed in the feces. Thus, the consequences of both bloodstream infections and pneumonia are that high levels of PA are excreted into the environment.

## DISCUSSION

PA is a threat to global health especially due to the increasing frequency of MDR infections. In particular, PA bloodstream infections, which are associated with mortality rates between 25 and 50%^2,21,22^, are an important clinical problem. Using *in vivo* imaging of bioluminescent bacteria and employing STAMP analysis, we identified an unexpected pathway through which PA traffics to the gallbladder and ultimately the intestines and feces. PA passes through narrow bottlenecks to gain access to the gallbladder, but this organ facilitates a dramatic expansion of PA numbers (Fig. 6). As a consequence, removal of the gallbladder dramatically reduces the levels of PA shed into the environment. In epidemiologic studies in humans and mice, the gallbladder has been shown to be an important replication niche for certain GI pathogens such as *Listeria monocytogenes* (LM), *Escherichia coli*, and *Salmonella enterica* serovar typhi^7,23,24^, and replication in the gallbladder has been shown to fuel transmission through gastrointestinal shedding. STAMP was previously applied to LM bloodstream infection revealing that LM experiences a similarly narrow bottleneck in the gallbladder and intestines^9^ and identifying the gallbladder as the main driver of fecal excretion. Here, we show for the first time that the same strategy is used by a *non-enteric* pathogen. Thus, migration of bacteria from the bloodstream to the gallbladder and subsequently to the intestines may be a more universal bacterial exit strategy than previously appreciated.

**Figure 6.**
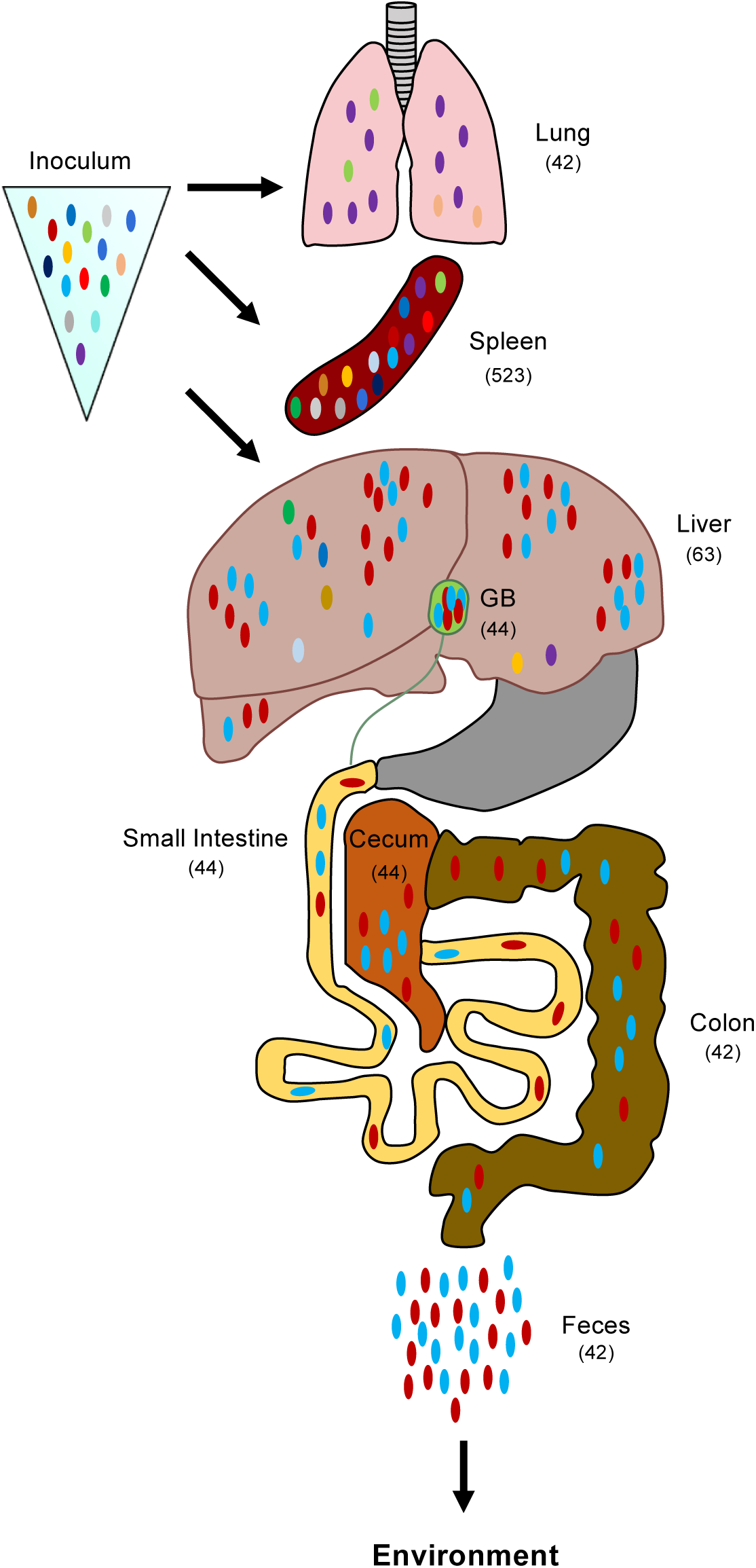
Model of *P. aeruginosa* population dynamics following bloodstream infection with barcoded organisms. *P. aeruginosa* populations with unique complexity arise in the lungs, liver, and spleen of intravenously infected animals. The bottleneck sizes at 24 hpi within the liver and lung are similar (mean *N*_b_′ values for all mice are shown in parentheses); however, their founding populations are distinct. Very few organisms establish infection in the gallbladder (GB), but the founders ultimately expand to very high numbers. Bacteria released from the GB through the bile into the small intestine become the principal source of *P. aeruginosa* excreted in the feces.

Systemic infections caused by opportunistic pathogens are often viewed as evolutionary and ecological “dead-ends” in that they are disadvantageous for pathogen survival and thought to be accidents that do not benefit the pathogen. Our work suggests that this paradigm does not apply to PA. Once in the bloodstream, PA transits to the gallbladder, which it exploits as a protected replication niche and uses to facilitate its own exit back into the environment via high levels of intestinal shedding. Mutations that enhance this process may therefore be selected for during infections and become fixed in the PA gene pool. Rather than being an ecological “dead-end,” PA bloodstream infections may instead serve as a transmission strategy (Fig. 6).

The hosts in which PA undergoes selection for gallbladder expansion and fecal excretion are less clear. PA naturally infects cows, dogs, cats, goats, and mink^25,26^, all of which have gallbladders^27^, so our mouse model may have identified bacterial processes that evolved in these mammals. Interestingly, several lines of evidence suggest that our findings may be applicable to humans. PA is one of the top 5 pathogens isolated from bile in patients with cholecystitis^28^ and has been associated with secondary sclerosing cholangitis, a dreaded complication of critical illness that involves the biliary system of the liver^29^. In this context, it has been suggested that PA gains access to the biliary tree by ascending from the duodenum via the common bile duct. However, it is conceivable that some of these infections are secondary to transient occult PA bacteremia and arrive in the gallbladder via the liver. Likewise, PA is a leading cause of acalculous cholecystitis (inflammation of the gallbladder in the absence of gallstones), a common complication of serious medical illness^30^, and it is similarly possible that this is the result of PA trafficking from the bloodstream to the gallbladder. Spread of PA to the intestines during systemic infections would have important clinical implications, as the GI tract is an important reservoir for PA in hospitalized patients^31–33^, and PA GI carriage is a precursor to both invasive infections^34,35^ and environmental contamination. Excreted PA is spread to other patients via the hands of healthcare workers^36–39^ or indirectly through contamination of objects such as bedrails, sinks, drains, and endoscopes^40,41^. Inter-institutional transfer of patients with GI carriage may facilitate the geographic spread of PA global epidemic strains such as sequence type (ST) 111, ST175 and ST235^42,43^. Our findings linking systemic PA infections with high levels of GI excretion via the gallbladder in mouse models support additional studies to examine the role of the gallbladder in PA transmission among patients.

Our findings suggest that PA is remarkably well equipped for replication in the gallbladder. Bile and bile salts have traditionally been viewed as toxic to bacteria^44^, but both laboratory-adapted and clinically-derived PA strains expanded in the gallbladders of mice, and select strains grew *ex vivo* in bile as well as in enriched medium. The mechanisms of bile resistance in PA are unknown; however, a role for multidrug efflux pumps such as MexAB-OprM has been suggested^45,46^. Thus, bile exposure may promote antibiotic resistance through increases in efflux pump expression, and transit through the gallbladder may enhance excretion of drug-resistant PA. Hardy and colleagues suggested that LM uses the lumen of the gallbladder to escape immune surveillance^23^, and it is possible that the immune-privileged nature of this organ provides a hospitable environment for PA replication as well. PA plays an active role in trafficking to and expansion in the gallbladder, as these processes are much less efficient in the absence of a functional T3SS. These findings suggest that, in addition to enhancing virulence, the PA T3SS promotes gastrointestinal carriage and transmission.

In summary, this work is the first to identify bacterial expansion in the gallbladder by the environmentally ubiquitous, non-enteric, opportunistic pathogen PA and suggests that the strategy of hijacking the gallbladder as a protected site for bacterial expansion to facilitate high levels of fecal excretion may be more widely utilized by bacterial species than previously thought. PA bacteremia leads to gallbladder expansion and fecal excretion, challenging the clinical paradigm that bloodstream infections are “dead-ends”. If these findings are shown to be applicable to humans, a better understanding of the gallbladder’s role in fecal excretion will be critical to curbing MDR bacterial transmission in the healthcare setting.

## METHODS

### Bacterial Strains and Growth Conditions

PA strains PABL002, PABL012, PABL016, PABL017, PABL028, PABL030, PABL041, PABL046, PABL049, PABL095, PABL107 are clinical isolates cultured from the bloodstreams of patients at Northwestern Memorial Hospital between September 1, 1999 and June 9, 2002^21^. PA14 and PAO1 are commonly used laboratory strains^47,48^. PAC6 and S10 are clinical isolates from the bloodstreams of patients in Taiwan with Shanghai Fever^16^. Relevant characteristics of these strains are listed in Table 1.

*Escherichia coli* strain TOP-10 (Invitrogen) was used for cloning and *E. coli* strains S17.1 λpir^49^ and SM10 λpir (gift of John Mekalanos, Harvard Medical School) were used to introduce plasmids into *P. aeruginosa*. Bacterial strains were streaked from frozen cultures onto either Vogel-Bonner minimal (VBM) agar^50^ or Luria Bertani (LB) agar and, unless otherwise stated, grown at 37°C in either LB or MINS media^51^.

Antibiotics were used at the following concentrations: irgasan 5 µg/mL (irg), gentamicin 100 μg/mL (gm), carbenicillin 250 μg/mL ^52^, and tetracycline 75 μg/mL (tet) for PA and gentamicin 15 μg/mL, carbenicillin 50 μg/mL, and tetracycline 10 μg/mL for *E. coli.* For blue-white discrimination, 5-bromo-4-Chloro-3-Indolyl-D-Galactopyranoside (Xgal) was added to plates at a concentration of 80 μg/mL. Further details on the strains and plasmids used in this study can be found in Tables 1, 2 and 3.

### Generation of Labeled *P. aeruginosa* Strains Expressing Luciferase, Gentamicin-resistance, or LacZ

The plasmid pminiCTXnpt2lux^53,54^, was transformed into *E. coli* strain S17.1 λpir. Following conjugation, the luciferase cassette was introduced into the chromosomal *attB* site of the PA strains PAO1, PA14, PABL012 and PABL012Δ*pscJ,* PABL002, PABL016, PABL017, PABL028, PABL030, PABL041, PABL046, PABL049, PABL095, PABL107, PAC6, and S10 via integrase-mediated recombination using previously defined approaches^55^ to generate luciferase-labeled strains (e.g. PABL012*_lux_*). Bioluminescent bacteria were confirmed by imaging plates on the IVIS Lumina LTE ® *in vivo* Imaging System.

The gentamicin resistance cassette, including its native promoter, was amplified from pEX18.Gm^56^ with the primers Gm-pCTX_F-BamHI-2 and Gm-pCTX_R-EcoRI (Table 4). It was then ligated into the multiple cloning site of pminiCTX-1^55^ to generate the plasmid pminiCTX-1*_GM_*. As above, the resulting plasmid was transformed into *E. coli* S17.1 λpir and, following conjugation, was introduced into PABL012 to generate the strain, PABL012*_GM_*. The gentamicin resistance cassette in PABL012*_GM_* was shown to have no impact on CFU recovery (data not shown).

The pminiCTX-1*_lacZ_* plasmid (gift from Stephen Lory, Harvard Medical School) was transformed into *E. coli* strain SM10 λpir. Following conjugation, the *lacZ* gene was introduced into the *attB* site of the strain PABL012 via integrase-mediated recombination using previously described approaches to generate the strain, PABL012*_lacZ_*.

### Generation of Type-3-Secretion-Deficient *P. aeruginosa*

Upstream and downstream fragments surrounding the *pscJ* gene were amplified from PAO1 genomic DNA using the following primers: pscJ-5-1-HindIII, pscJ-5-2, pscJ-3-1, and pscJ-3-2 HindIII, where pscJ-5-2 and pscJ-3-1 contain a 24 bp-overlapping linker sequence (*TTCAGCATGCTTGCGGCTCGAGTT*) to generate an in-frame deletion of the *pscJ* gene (Table 4). The integration proficient vector, pEXG2^57^ was cut with HindIII, and the three fragments were ligated using the New England Biolabs Gibson Assembly ® Cloning Kit. The resulting vector, pEXG2Δ*pscJ*, was verified by sequencing at the NuSeq facility at Northwestern University and transformed into *E. coli* SM10 λpir. Following conjugation and allelic exchange with PABL012*_lux_*, whole-genome sequencing was performed on strain PABL012Δ*pscJ_lux_* to confirm the mutation.

### Mouse Model of Tail Vein Injection

Overnight cultures of PA were grown in 5 mL of MINS medium, diluted into fresh medium the next day, and regrown to exponential phase prior to infections. The tail veins of female 6- to 8-week-old BALB/c mice restrained using a TV-150 Tailveiner ® (Braintree Scientific) were dilated with a heat lamp. A defined number of PA bacteria in 50 µL of phosphate buffered saline (PBS) was injected into the tail veins of mice using a 27-gauge needle. Inoculums were confirmed by plating serial dilutions on LB agar. To monitor bacterial shedding, fecal pellets were harvested individually from each mouse, weighed, and re-suspended in PBS. At specified times post-infection, the mice were anesthetized and sacrificed by cervical dislocation. For dissemination experiments, organs were excised, weighed, and homogenized in PBS using the Benchmark ® Bead Blaster 24. Viable bacteria were enumerated by plating serial dilutions of either fecal pellets or organs on LB agar containing 5 μg/mL of irgasan to select for PA. Luminescent bacteria were confirmed by imaging plates on the IVIS Lumina LTE ® *in vivo* Imaging System. Recovered bacteria were counted to determine CFU per gram organ.

Animals were purchased from Harlan Laboratories, Inc. (BALB/c, Indianapolis, IN) and Jackson Laboratory (C57/BL6, Bar Harbor, ME) and housed in the containment ward of the Center for Comparative Medicine at Northwestern University. All experiments were approved by the Northwestern University Animal Care and Use Committee.

### Mouse Model of Oral Gavage

Briefly, overnight cultures of PA were grown in 5 mL of MINS medium, diluted into fresh medium the next day, and regrown to exponential phase prior to infections. Mice were restrained, and a 20-gauge blunt-end straight feeding needle (Pet Surgical, Sherman Oaks, CA) was used to gavage 50 µL of the appropriate dose of PA directly into the stomachs of 6- to 8-week-old BALB/c mice. Again, inoculums were confirmed by plating serial dilutions on LB agar. Organ harvesting and bacterial plating for CFU enumeration was performed as described above.

### Mouse Model of Acute Pneumonia

The mouse model of acute pneumonia described by Comolli, et al.^58^ was used for all animal experiments. Briefly, 6- to 8-week-old female BALB/c mice were anesthetized by intraperitoneal injection of a mixture of ketamine (75 mg/kg) and xylazine (5 mg/kg). Mice were intranasally inoculated with specified doses of bacteria in 50 μL of PBS. Inoculums were confirmed by plating serial dilutions on LB agar. Organ harvesting and bacterial plating for CFU enumeration was performed as described above.

### Mouse Cholecystectomy and Sham Surgery

Briefly, 4- to 6-week-old BALB/c mice were transported to the Northwestern Microsurgery Core at Northwestern University. There, mice were anaesthetized with single doses of ketamine (100 mg/kg) and xylazine (20 mg/kg) and administered a subcutaneous one-time dose of 0.05 mg/kg buprenorphine prior to incision. Nair was used to remove abdominal hair and the mouse was positioned on its dorsal surface under an operating microscope. A one-inch skin incision was made in the abdominal skin followed by a similar incision in the peritoneum. The peritoneum was reflected and the sternum retracted. Bowels were covered in sterile gauze infused with warm saline. The ligament attaching the murine gallbladder to the base of the diaphragm was severed with electrocautery. The gallbladder was then tied off at the base with an absorbable 7-0 silk suture, excised in total with electrocautery. The abdominal cavity was irrigated with warm saline and the peritoneum closed with a 5-0 absorbable suture. The abdominal skin was closed with a 5-0 nylon, non-absorbable suture using a running stitch. Mice were given an immediate post-operative dose of subcutaneous Meloxicam (1 mg/kg) and recovered at 32°C for 24 hours. At 24 hours post operation, mice were given an additional one-time subcutaneous dose of 1 mg/kg of Meloxicam. For ‘sham’ mice, the operative protocol was identical to above with the exception that ligaments were not severed and gallbladders were not removed. Mice recovered for one week post-operatively prior to challenge with PA in the containment ward of the Center for Comparative Medicine at Northwestern University. During this phase, mice were monitored daily for signs of distress and infection. All experiments were approved by the Northwestern University Animal Care and Use Committee.

### Analysis of *in vivo* Localization of Bacteria Using IVIS Imaging System

Representative 6- to 8-week-old female BALB/c mice were challenged either intravascularly or intranasally with various bioluminescent PA strains. At specific times post-infection, mice were anesthetized with vaporized isoflurane using the XGI-8 Gas Anesthesia System, transferred to the stage of the IVIS Lumina LTE ® *in vivo* Imaging System, and both photographic and bioluminescent images were captured. All images were captured with an exposure time of two minutes and medium binning. Bioluminescence is represented as a heat map (red = most intense) normalized to a range of 1 x 10^4^ to 1 x 10^5^ radiance (p/sec/cm^2^/sr). Images were processed with Living Image ® Software version v4.0 by Caliper Life Sciences.

### Construction of a Library of Barcoded *P. aeruginosa*

The STAMP protocol used here was similar to that described by Abel et al. and Zhang et al.^9,11^. Table 4 contains the sequences of all primers used in this protocol. The plasmid pminiCTX*_STAMP_* used for generating the tagged *P. aeruginosa* library was constructed as follows. The gentamicin resistance cassette (∼1032 bp) from pEX18.Gm^56^ was amplified with primers P80 and P110^11^, the latter of which is degenerate and contains a string of 30 random bases (Table 4). The amplified product was inserted into the EcoRI site of the integration proficient plasmid pminiCTX-1^55^. The resulting plasmid, pminiCTX*_STAMP_*, was transformed into *E. coli* strain TOP-10 and selected with gentamicin. Plasmid DNA was harvested and transformed into *E. coli* SM10 λpir. Following conjugation, pminiCTX*_STAMP_* was introduced into the *attB* site of the PA strain PABL012 via integrase-mediated recombination to generate PABL012*_pool_*, and transconjugants were selected with tetracycline. After 10 of 10 tested colonies were found to have a unique, correct insertion of pminiCTX*_STAMP_*, the remaining colonies were scraped off plates and pooled. After addition of 25% glycerol, the pooled library of PABL012 (designated “PABL012*_pool_*”) was divided into aliquots and stored at −80°C. The plasmid was stably integrated for at least 20 hours without selection (Extended Data Fig. 5B).

### Calibration Curve for the STAMP Study

As described by Abel et al.^11^, the diversity of barcodes present in a subculture of tagged bacteria can be used to calculate the size of the founding population of that subculture. We compared the mathematically determined founding population sizes to those induced experimentally. Briefly, we diluted three frozen aliquots of the PABL012*_pool_* library (A, B, C) (50 μL) 1:100 in MINS and grew with shaking overnight at 37°C. In the morning, cultures were diluted into fresh MINS medium and grown at 37°C for 3 hours. Cells were harvested by centrifugation (13,000 x g, room temperature, 2 min) and resuspended in PBS. A small portion of each serial dilution of each replicate was used to enumerate total CFU, and the remainder was plated at 37°C overnight. Following growth, all bacterial colonies were scraped off using Falcon cell scrapers into 5 mL of PBS. Samples were concentrated by centrifugation at 8,000 x g for 10 minutes and resuspended in 1 mL of PBS. Genomic DNA was harvested from each sample using the Promega Maxwell 16 Instrument ® and the Cell DNA Purification Kit. The tagged region that harbored the 30-bp barcode was amplified in triplicate from genomic DNA using primer P47 and primers P48, P51-73 (Table 4). The PCR products were run on a 1% agarose gel, pooled, extracted from the gel using Qiagen QIAquick ® gel extraction kit and quantified (Invitrogen Quant-iT^™^ dsDNA Assay Kit, High Sensitivity). The purified PCR products were combined in equimolar concentrations and sequenced on an Illumina Miseq instrument (Miseq Reagent Kit V2, 50-cycle, Illumina) using custom sequencing primer P49^11^ (Table 4) with a mean cluster density of 8.6 x 10^5^ ± 2.5 x 10^5^. Reaper-15-065 was used to discard sequence reads with low quality (≤Q30) and to trim the sequence following the barcode^59^. The trimmed sequences were clustered with QIIME (version 1.9.1) using pick_otus.py with a sequence similarity threshold of 0.9^60,61^. The resulting estimated founding population sizes (*N_b_*) were mathematically calculated from the frequency of each barcode as described by Abel et al.^11^ using the method of Krimbas and Tsakas^62^. These *N_b_* values were then compared to experimentally determined founding population sizes (CFU) to generate a calibration curve (Extended Data Fig. 4C). As mathematically derived *N_b_* values underestimated the actual CFU values, this calibration curve was used to generate adjusted founding population sizes (*N_b_*′) by correcting *N_b_* values obtained from mouse experiments for this underestimation. For each *N_b_*′ value calculated for every organ in each mouse, 95% confidence intervals are presented (Extended Data Figure 5D, 6D).

### Determination of barcode distribution, allelic frequency, and skew present in PABL012*_pool_*

Barcodes from 30 independently sequenced technical replicates from the same PABL012*_pool_* inoculum sample were analyzed as described above. Individual barcodes present in at least 29 of 30 aliquots were deemed adequately represented in the PABL012*_pool_* inoculum given to mice and were included in the calculation of *N_b_* values^11^ (14 hpi n=4612, 24 hpi n=4208, INOC30). Barcodes not present in at least 29 of 30 aliquots were filtered from the barcodes recovered from *in vivo* experiments. To assess input barcode distribution, the maximum allelic frequency of all barcodes present in INOC30 was arrayed on the x-axis in descending frequency (Extended Data Fig. 5B, 6B), which revealed that the maximum allelic frequency of any single barcode in the total population at both 14 and 24 hpi was 1.2%. The vast majority of barcodes (>99%) were present in the population at <0.5% (allelic frequency <0.005). To assess the impact of this minor input barcode (inoculum) skew on the output barcode (mouse organ) recovery, the maximum input frequency of each barcode present in INOC30 was plotted on the y-axis verses the maximum output barcode frequency of the identical barcode recovered from any mouse organ (Extended Data Figure 5C, 6C). Linear regression with best fit analysis was performed and yielded ‘goodness of fit’ R_2_ values of 0.157 and 0.097, respectively, indicating that input barcode allelic frequency had a poor correlation to output barcode frequency and demonstrating the stochastic nature of barcode loss in our model (Graph Pad Prism v7.0b). Had the allelic frequencies of the input barcodes dictated the output allelic frequencies, we would anticipate an observed an R_2_ value closer to 1.

### Animal Infections with PABL012*_pool_*

An inoculum of barcoded PABL012*_pool_* was prepared and injected into mice as described above. At 14 and 24 hpi, mice were euthanized, and the lungs, liver, spleen, gallbladder, stomach, small intestine, cecum, colon, and feces were harvested, weighed, homogenized in 1 mL of PBS, and plated for CFU counts. For *N_b_*′ analyses, 500 µL of samples from the lung, spleen, liver, gallbladder, stomach, small intestine, cecum, colon, feces, and 250 µL of inoculum x 30 were independently spread on 100 x 15 mm Petri dishes containing LB with 5 µg/mL irgasan. Plates were grown at 37°C overnight and bacteria sequenced for barcode diversity as above. *N_b_* values were calculated as described in the previous section and adjusted using the calibration curve with an R script^11^ to estimate *in vivo* founding population sizes (*N_b_*′). When the number of recovered CFUs was below 100, *N_b_*′ determination was not performed. “Genetic distance” between the bacterial populations in two different organs was estimated by comparing the barcode allelic frequencies of the two populations using the Cavalli-Sforza chord distance method^13^ as described by Abel et al.^11^. Genetic relatedness (GR) was calculated as (1 – genetic distance).

### Genetic Relatedness (GR) Threshold Determination

Given that the 30 inoculum replicates (INOC30) were from the same original sample of PABL012*_pool_* and therefore technically identical with the maximum possible experimentally determined GR, we averaged the GR between inoculum samples at 14 and 24 hpi. We then used this value minus 2 standard deviations to set the lower threshold for “high” levels of GR at ≥ 0.88. Given that barcode loss is stochastic and that the dominant barcodes found in individual organs are unique to each mouse, we determined inter-mouse (between mice) and intra-mouse (within a single mouse) GR between organs at 14 and 24 hpi (Extended Data Fig. 4D, Figure 2B,D). We averaged all the inter-mouse and intra-mouse GR values, which yielded a value of 0.41. This value was used to set the lower threshold for “moderately high” GR (i.e. “moderately high” is GR values ≥ 0.41 and ≤ 0.88). Since populations of PA in organs between mice (inter-mouse) were the least likely to be related, we averaged the GR values of bacteria from these organs (inter-mouse LG, LV, ST, GB, INT, CM, CO, FE at 14 and 24 hpi) to set the upper threshold of the “low” GR category as ≤ 0.17. The inter-mouse spleen GR values were specifically excluded from this calculation because they displayed moderately high (14 hpi) and moderately low (24hpi) GR respectively, likely secondary to the presence of large numbers of diverse barcodes in this organ (Extended Data Fig. 4D).

### Electron Microscopy

For transmission electron microscopy (TEM), 6- to 8-week-old female BALB/c mice were infected via tail vein as described above with either PABL012*_lux_* or PBS. At defined timepoints post-injection, gallbladders were harvested, fixed for at least 48 hours at 4°C in 0.1 M sodium cacodylate buffer (pH 7.3) containing 2% paraformaldehyde and 2.5% glutaraldehyde, and post-fixed with 2% osmium tetroxide in unbuffered aqueous solution. Samples were rinsed with distilled water, *en bloc* stained with 3% uranyl acetate, and rinsed again with distilled water. They were then dehydrated in ascending grades of ethanol, transitioned with propylene oxide, embedded in the resin mixture of Embed 812 kit, and cured in a 60°C oven. Samples were sectioned on a Leica Ultracut UC6 ultramicrotome. One-micrometer thick sections were collected and stained with Toluidine Blue O. Seventy-nanometer (thin) sections were collected on 200 mesh copper grids and stained with uranyl acetate and Reynolds lead citrate. Images were obtained using the FEI Tecnai Spirit G2 transmission electron microscope with the help of the Northwestern Center for Advanced Microscopy (CAM).

### Histology

For histological sections, 6- to 8-week-old female BALB/c mice were injected via tail vein as described above with either PABL012*_lux_*, PABL012Δ*pscJ_lux_*, or PBS. At defined points post-injection, gallbladders were harvested and fixed in 4% paraformaldehyde solution for a minimum of 48 hours. Samples were then paraffin embedded, processed, and stained with hematoxylin and eosin (H&E) by the Mouse Histology and Phenotyping Laboratory (MHPL) at Northwestern University. Gallbladder sections were imaged on the Zeiss Axioscope with a CRI Nuance Camera, located in the Northwestern University CAM.

### Bile Growth Assays

*P. aeruginosa* PABL012*_lux_* or PAO1*_lux_* were grown overnight in MINS medium, inoculated 1:8 into fresh medium the following day and regrown to exponential phase. One milliliter of culture was pelleted, re-suspended in PBS, and the OD_600_ adjusted to 0.7. This culture was diluted 1:100 and inoculated in triplicate into 180 µL of LB, PBS, MINS medium, or a 35-40% bile solution. To prepare the bile solution, 6- to 12-week-old BALB/c mice were anesthetized and sacrificed by cervical dislocation. Gallbladders were harvested intact and lanced to release bile contents. Gentle centrifugation was used to pellet tissue contents, liberating free bile. Freshly harvested bile was diluted to 35-40% with PBS. Following inoculation, cultures were grown with shaking at 37°C for 24 hours. Samples were taken at time 0, 4, and 24 hours, diluted, and plated to quantify CFU.

### Minimal Inhibitory Concentrations

Minimal inhibitor concentration (MICs) for all PA strains used were determined in triplicate using the broth microdilution protocol described by Wiegand, et al.^63^ (Table 1). The following antibiotics were prepared from commercially available sources and were used to assess MICs: piperacillin/tazobactam (Pip/Tazo), cefepime (Cep), ceftazidime (Ctz), ciprofloxacin (Cipro), meropenem (Mero), gentamicin (Gent), colistin (Col), and aztreonam (Az).

### Statistical Analysis and Data Availability

Analyses were performed with the help of Graph Pad Prism v7.0b. Data are represented as geometric mean (horizontal line or box) +/- standard deviation when multiple mice were challenged. Bacterial CFU is represented as CFU per gram organ and organs from individual mice are represented by a single data point on each graph (circle if CFU recovered, diamond if no CFU recovered). A non-parametric ANOVA test with Kruskal-Wallis test correction for multiple comparisons was used to compare bacterial growth in specific media given small samples sizes and non-normal distributions. The data that support the findings of this study are available from the corresponding author upon request.

## SUPPLEMENTAL DATA

Table 1. P. *aeruginosa* Strain Summary.

Table 2. Bacterial strains used in this study.

Table 3. Plasmids used in this study.

Table 4. Primers used in this study.

Extended Data Figure 1. Bile promotes the replication of *P. aeruginosa*.

Extended Data Figure 2. Gastrointestinal shedding of *P. aeruginosa* appears universal and independent of mouse genetics or gender.

Extended Data Figure 3. Dissemination to the gallbladder and fecal sheeding are shard features among *exoS*- and *exoU*-containing clinical isolates of *P. aeruginosa*.

Extended Data Figure 4. STAMP control experiments and calculations.

Extended Data Figure 5. Analysis of individual STAMP barcode distributions across all organs in all mice at 14 hpi.

Extended Data Figure 6. Analysis of individual STAMP barcode distributions across all organs in all mice at 24 hpi.

## ACKNOWLEDGEMENTS

We thank S. Abel and P. Abel for providing the scripts, analysis pipeline, and discussion regarding STAMP, S. Lory, J. Mekalanos, and C.-H. Chiu for providing *P. aeruginosa* strains and cloning tools, L. Reynolds for technical assistance with electron microscopy, and S. Han and J. Zhang for mouse surgery. This work was supported by grants from the National Institutes of Health/National Institute of Allergy and Infectious Disease (R01 AI118257, R01 AI053674, U19 AI135964, K24 104831, and R21 AI129167, awarded to A.R.H.) an American Cancer Society Postdoctoral Fellowship (130602-PF-17-107-01-MPC, awarded to K.E.R.B.), and an American Heart Association Postdoctoral Fellowship (73POST25830019, awarded to J.P.A). Additional support was provided by the Northwestern University Mouse Histology and Phenotyping Laboratory supported by an NCI Cancer Center Support Grant (CCSG NCI P30-CA060553). Imaging work was performed at the Northwestern Center for Advanced Microscopy (CAM) also generously supported by NCI CCSG P30 CA060553. Mouse surgery was performed by the Northwestern University Comprehensive Transplant Center (NUCTC) Microsurgery Core. The funders had no role in study design, data collection and analysis, decision to publish, or preparation of the manuscript. We would like to thank all members of the Hauser, Ozer, and Kociolek laboratories for their valuable comments during numerous discussions of this work.

## AUTHOR CONTRIBUTIONS

K.E.R.B. and J.P.A. designed and performed the experiments, analyzed and interpreted the data, and wrote the manuscript. B.H.C. performed experiments and C.-H. C. provided intellectual discussion and wrote the manuscript. A.R.H. provided resources and intellectual contributions, oversaw interpretation, and wrote the manuscript. All authors discussed the results and commented on the manuscript.

## CORRESPONDENCE AND MATERIALS REQUESTS

Should be addressed to K.E.R.B. at 303 E. Chicago Ave., Ward 6-035, Chicago, IL 60611. Phone: (312) 503-1081. Fax: (312) 503-1339.

