## Supplemental Tables and Figures for "Systemic Infection Facilitates Transmission of *Pseudomonas aeruginosa*"

Table 1. *P. aeruginosa* Strain Summary

| | | | | | | MICs (µg/mL) | | | | | | | |
|---|---|---|---|---|---|---|---|---|---|---|---|---|---|
| **Strain** | **ST Type** | **Source** | **Geography** | **NCBI Accession Number** | ***exoU/S*** | **Pip/Tazo** | **Cep** | **Ctz** | **Mero** | **Cipro** | **Gent** | **Col** | **Az** |
| PABL002 | 253 | Bloodstream | Chicago | GCA_003412285.1 | U | **2/4** | **1** | **2** | **<0.25** | **<0.25** | **1** | **1** | **4** |
| PABL012 | Unk. | Bloodstream | Chicago | CP031659.1 | S | **4/4** | **2** | **2** | **0.5** | **<0.25** | **1** | **1** | **4** |
| PABL016 | 2555 | Bloodstream | Chicago | GCA_003412165.1 | U | **4/4** | **2** | **1** | **<0.25** | **<0.25** | **2** | **4** | **4** |
| PABL017 | 2167 | Bloodstream | Chicago | CP031660.1 | S | **4/4** | **2** | **2** | **0.5** | **<0.25** | **2** | **1** | **4** |
| PABL028 | Unk. | Bloodstream | Chicago | GCA_003411845.1 | S | **4/4** | **2** | **2** | **<0.25** | **<0.25** | **4** | **1** | **4** |
| PABL030 | Unk. | Bloodstream | Chicago | GCA_003412115.1 | S | **4/4** | **1** | **1** | **0.5** | **<0.25** | **2** | **1** | **8** |
| PABL041 | 253 | Bloodstream | Chicago | GCA_003411465.1 | U | **4/4** | **1** | **1** | **1** | **<0.25** | **1** | **1** | **4** |
| PABL046 | Unk. | Bloodstream | Chicago | GCA_003411535.1 | S | **16/4** | **8** | **2** | **8** | **128** | **8** | **1** | **16** |
| PABL049 | 244 | Bloodstream | Chicago | GCA_003411745.1 | S | **8/4** | **4** | **2** | **4** | **<0.25** | **1** | **2** | **16** |
| PABL095 | 348 | Bloodstream | Chicago | GCA_003410805.1 | S | **8/4** | **4** | **2** | **4** | **4** | **1** | **2** | **8** |
| PABL107 | 377 | Bloodstream | Chicago | GCA_003410475.1 | U | **8/4** | **4** | **2** | **4** | **<0.25** | **1** | **2** | **4** |
| PAC6 | 1021 | Bloodstream | Taiwan | | U | **8/4** | **1** | **2** | **1** | **<0.25** | **1** | **1** | **8** |
| S10 | 244 | Bloodstream | Taiwan | | S | **8/4** | **1** | **1** | **1** | **<0.25** | **1** | **1** | **8** |
| PA01 | 549 | Wound | Australia | NC_002516.2 | S | **4/4** | **2** | **2** | **2** | **<0.25** | **2** | **2** | **8** |
| PA14 | 253 | Wound | Unk. | NC_008463.1 | U | **4/4** | **2** | **2** | **2** | **<0.25** | **1** | **1** | **8** |

U = exoenzyme U, S = exoenzyme S; Unk. = unknown
Pip/Tazo = piperacillin-tazobactam, Cep = cefepime, Ctz = ceftazidime, Mero = meropenem, Cipro = ciprofloxacin, Gent = gentamicin, Col = Colistin, Az = aztreonam
Resistant, Intermediate: Clinical Laboratory Standards Institute, MIC Interpretive Standards (µg/mL), 2018

Table 2. Bacterial strain list

| Species | Strain ID | Relevant Characteristics | Reference |
|---|---|---|---|
| *E. coli* | TOP-10 | F- mcrA Δ(mrr-hsdRMS-mcrBC) φ80lacZΔM15 ΔlacX74 nupG recA1 araD139 Δ(ara-leu)7697 galE15 galK rpsL(StrR) endA1 λ- | Invitrogen |
| *E. coli* | S17-1 λpir | $Sm_R$; *pro, thi, hsd*$R_-M_+$, RP4-2-Tc:Mu;Km:Tn7 λpir | Simon (1983) |
| *E. coli* | SM10 λpir | $Km_R$*; thi*-1 *thr, leu, tonA, lacY, supE, recA*::RP4-2-Tc::Mu λpir | Simon (1983) |
| *P. aeruginosa* | PABL002 | human bacteremia isolate (presumed source: intraabdominal abscess) | Scheetz (2009) |
| *P. aeruginosa* | $PABL002_{lux}$ | As above, luciferase cassette cloned into attB site | This study |
| *P. aeruginosa* | PABL012 | human bacteremia isolate (presumed source: uti) | Scheetz (2009) |
| *P. aeruginosa* | $PABL012_{lux}$ | As above, luciferase cassette cloned into attB site | This study |
| *P. aeruginosa* | $PABL012_{GM}$ | As above, with gentamicin cassette cloned into attB site | This study |
| *P. aeruginosa* | $PABL012_{lacZ}$ | As above, with *lacZ* cloned into attB site | This study |
| *P. aeruginosa* | PABL012Δ*pscJ* | Clean deletion of *pscJ* (Δ5-245) rendering strain T3S deficient | This study |
| *P. aeruginosa* | PABL012Δ$pscJ_{lux}$ | As above, with luciferase cassette cloned into attB site | This study |
| *P. aeruginosa* | PABL016 | human bacteremia isolate (presumed source: Indwelling catheter) | Scheetz (2009) |
| *P. aeruginosa* | $PABL016_{lux}$ | As above, luciferase cassette cloned into attB site | This study |
| *P. aeruginosa* | PABL017 | human bacteremia isolate (presumed source: pneumonia) | Scheetz (2009) |
| *P. aeruginosa* | $PABL017_{lux}$ | As above, luciferase cassette cloned into attB site | This study |
| *P. aeruginosa* | PABL028 | human bacteremia isolate (presumed source: pneumonia) | Scheetz (2009) |
| *P. aeruginosa* | $PABL028_{lux}$ | As above, luciferase cassette cloned into attB site | This study |
| *P. aeruginosa* | PABL030 | human bacteremia isolate (presumed source: unknown) | Scheetz (2009) |
| *P. aeruginosa* | $PABL030_{lux}$ | As above, luciferase cassette cloned into attB site | This study |
| *P. aeruginosa* | PABL041 | human bacteremia isolate (presumed source: unknown) | Scheetz (2009) |
| *P. aeruginosa* | $PABL041_{lux}$ | As above, luciferase cassette cloned into attB site | This study |
| *P. aeruginosa* | PABL046 | human bacteremia isolate (presumed source: unknown) | Scheetz (2009) |
| *P. aeruginosa* | $PABL046_{lux}$ | As above, luciferase cassette cloned into attB site | This study |
| *P. aeruginosa* | PABL049 | human bacteremia isolate (presumed source: biliary stent) | Scheetz (2009) |
| *P. aeruginosa* | $PABL049_{lux}$ | As above, luciferase cassette cloned into attB site | This study |
| *P. aeruginosa* | PABL095 | human bacteremia isolate (presumed source: pneumonia) | Scheetz (2009) |
| *P. aeruginosa* | $PABL095_{lux}$ | As above, luciferase cassette cloned into attB site | This study |
| *P. aeruginosa* | PABL107 | human bacteremia isolate (presumed source: unknown) | Scheetz (2009) |
| *P. aeruginosa* | $PABL107_{lux}$ | As above, luciferase cassette cloned into attB site | This study |
| *P. aeruginosa* | PAC6 | human bacteremia isolate (Shanghai Fever) | Chuang (2014) |
| *P. aeruginosa* | $PAC6_{lux}$ | As above, luciferase cassette cloned into attB site | This study |
| *P. aeruginosa* | S10 | human bacteremia isolate (Shanghai Fever) | Chuang (2014) |
| *P. aeruginosa* | $S10_{lux}$ | As above, luciferase cassette cloned into attB site | This study |
| *P. aeruginosa* | PAO1 | human wound isolate | Halloway (1955) |
| *P. aeruginosa* | $PAO1_{lux}$ | As above, luciferase cassette cloned into attB site | This study |
| *P. aeruginosa* | PA14 | human wound isolate | Rahme (1995) |
| *P. aeruginosa* | $PA14_{lux}$ | As above, luciferase cassette cloned into attB site | This Study |

Table 3. Plasmid list

| Plasmid | Relevant Characteristics | Reference |
|---|---|---|
| pEXG2 | Gm$_R$; allelic exchange vector with pBR origin, *sacB*+ | Rietsch (2005) |
| pEXG2-Δ*pscJ* | Gm$_R$; allelic exchange vector for making unmarked deletion of *pscJ* (Δ5-245); strains are null for T3 secretion | This study |
| pFLP2 | Ap$_R$/Cb$_R$; *oriT*+ *sacB*+, Flp recombinase | Hoang (1998) |
| pEX18.Gm | Gm$_R$; *oriT*+ *sacB*+, gene replacement vector with MCS from pUC18 | Hoang (1998) |
| pminiCTX-1 | Tc$_R$; self-proficient integration vector with Ω-*FRT*-*attP*-MCS, *ori, int,* and *oriT* | Hoang (2000) |
| pminiCTX*npt2lux* | Tc$_R$; luciferase expressing cassette driven by *pnpt2* promoter ligated into miniCTX-1 | Diaz (2008) |
| pminiCTX-1$_{GM}$ | Tc$_R$ Gm$_R$; Gentamicin resistance cassette (amplified from pEX18Gm) driven by native promoter ligated into miniCTX-1 | This study |
| pminiCTX-1$_{lacZ}$ | Tc$_R$; *lacZ* expressing cassette ligated into miniCTX-1 | Gift of S. Lory |
| pminiCTX$_{STAMP}$ | Tc$_R$ Gm$_R$; barcoded Gentamicin resistance cassette (amplified from pEX18Gm) driven by native promoter ligated into EcoRI site in miniCTX-1 | This study |

Table 4. Primer list

| Primer | Sequence 5’-3’ |
|---|---|
| pscJ – 5-1 HindIII | GCATAAATGTAAAGCAGCAGCAGCGATCCTGC |
| pscJ - 5-2 | *AACTCGAGCCGCAAGCATGCTGAA*CGTTCGCCTCATTGGGTC |
| pscJ - 3 -1 | *TTCAGCATGCTTGCGGCTCGAGTT*CGGCAACGGGGCTGAGCG |
| pscJ - 3 -2 HindIII | CTAGAGTCGACCTGCAGACTGCTCCTGGTAAACCTCG |
| pscJ - 5 | CCATGGCAATCCCTGGCCGGC |
| pscJ - 3 | CCGGTAGTAGCGATTGCAGCGC |
| Gm-pCTX_F-BamHI-2 | CGGGATCCCGCCGCTCATGAGACAATAACCCTGA |
| Gm-pCTX_R-EcoRI | CGGAATTCCGAAAGTATATATGAGTAAACTT |
| P49 | ACGCTCTTCCGATCTTGTAAAACGACGGCCAGT |
| P47 | AATGATACGGCGACCACCGAGATCTACACTCTTTCCCTACACGACGCTCTTCCGATCT |
| P80 | TATCGATAAGCTTGATATCGGTAAACTTGGTCTGACAATCGATGC |
| P110 | TGGATCCCCCGGGCTGCAGGCAGGAAACAGCTATGACNNNNNNNNNNNNNNNNNNNNNNNNNNNNNNNNNACTGGCCGTCGTTTTACACGCCGCTCATGAGACAATAACCCTGA |
| P48 | CAAGCAGAAGACGGCATACGAGATCGTGATGTGACTGGAGTTCAGACGTGTGCTCTTCCGATCATTCAGCAGACCTACGATGTCGGGG |
| P51 | CAAGCAGAAGACGGCATACGAGATACATCGGTGACTGGAGTTCAGACGTGTGCTCTTCCGATCATTCAGCAGACCTACGATGTCGGGG |
| P52 | CAAGCAGAAGACGGCATACGAGATGCCTAAGTGACTGGAGTTCAGACGTGTGCTCTTCCGATCATTCAGCAGACCTACGATGTCGGGG |
| P53 | CAAGCAGAAGACGGCATACGAGATTGGTCAGTGACTGGAGTTCAGACGTGTGCTCTTCCGATCATTCAGCAGACCTACGATGTCGGGG |
| P54 | CAAGCAGAAGACGGCATACGAGATCACTGTGTGACTGGAGTTCAGACGTGTGCTCTTCCGATCATTCAGCAGACCTACGATGTCGGGG |
| P55 | CAAGCAGAAGACGGCATACGAGATATTGGCGTGACTGGAGTTCAGACGTGTGCTCTTCCGATCATTCAGCAGACCTACGATGTCGGGG |
| P56 | CAAGCAGAAGACGGCATACGAGATGATCTGGTGACTGGAGTTCAGACGTGTGCTCTTCCGATCATTCAGCAGACCTACGATGTCGGGG |
| P57 | CAAGCAGAAGACGGCATACGAGATTCAAGTGTGACTGGAGTTCAGACGTGTGCTCTTCCGATCATTCAGCAGACCTACGATGTCGGGG |
| P58 | CAAGCAGAAGACGGCATACGAGATCTGATCGTGACTGGAGTTCAGACGTGTGCTCTTCCGATCATTCAGCAGACCTACGATGTCGGGG |
| P59 | CAAGCAGAAGACGGCATACGAGATAAGCTAGTGACTGGAGTTCAGACGTGTGCTCTTCCGATCATTCAGCAGACCTACGATGTCGGGG |
| P60 | CAAGCAGAAGACGGCATACGAGATGTAGCCGTGACTGGAGTTCAGACGTGTGCTCTTCCGATCATTCAGCAGACCTACGATGTCGGGG |
| P61 | CAAGCAGAAGACGGCATACGAGATTACAAGGTGACTGGAGTTCAGACGTGTGCTCTTCCGATCATTCAGCAGACCTACGATGTCGGGG |
| P62 | CAAGCAGAAGACGGCATACGAGATTGTTGACTGTGACTGGAGTTCAGACGTGTGCTCTTCCGATCATTCAGCAGACCTACGATGTCGGGG |
| P63 | CAAGCAGAAGACGGCATACGAGATACGGAACTGTGACTGGAGTTCAGACGTGTGCTCTTCCGATCATTCAGCAGACCTACGATGTCGGGG |
| P64 | CAAGCAGAAGACGGCATACGAGATTCTGACATGTGACTGGAGTTCAGACGTGTGCTCTTCCGATCATTCAGCAGACCTACGATGTCGGGG |
| P65 | CAAGCAGAAGACGGCATACGAGATCGGGACGGGTGACTGGAGTTCAGACGTGTGCTCTTCCGATCATTCAGCAGACCTACGATGTCGGGG |
| P66 | CAAGCAGAAGACGGCATACGAGATGTGCGGACGTGACTGGAGTTCAGACGTGTGCTCTTCCGATCATTCAGCAGACCTACGATGTCGGGG |
| P67 | CAAGCAGAAGACGGCATACGAGATCGTTTCACGTGACTGGAGTTCAGACGTGTGCTCTTCCGATCATTCAGCAGACCTACGATGTCGGGG |
| P68 | CAAGCAGAAGACGGCATACGAGATAAGGCCACGTGACTGGAGTTCAGACGTGTGCTCTTCCGATCATTCAGCAGACCTACGATGTCGGGG |
| P69 | CAAGCAGAAGACGGCATACGAGATTCCGAAACGTGACTGGAGTTCAGACGTGTGCTCTTCCGATCATTCAGCAGACCTACGATGTCGGGG |
| P70 | CAAGCAGAAGACGGCATACGAGATTACGTACGGTGACTGGAGTTCAGACGTGTGCTCTTCCGATCATTCAGCAGACCTACGATGTCGGGG |
| P71 | CAAGCAGAAGACGGCATACGAGATATCCACTCGTGACTGGAGTTCAGACGTGTGCTCTTCCGATCATTCAGCAGACCTACGATGTCGGGG |
| P72 | CAAGCAGAAGACGGCATACGAGATATATCAGTGTGACTGGAGTTCAGACGTGTGCTCTTCCGATCATTCAGCAGACCTACGATGTCGGGG |
| P73 | CAAGCAGAAGACGGCATACGAGATAAAGGAATGTGACTGGAGTTCAGACGTGTGCTCTTCCGATCATTCAGCAGACCTACGATGTCGGGG |

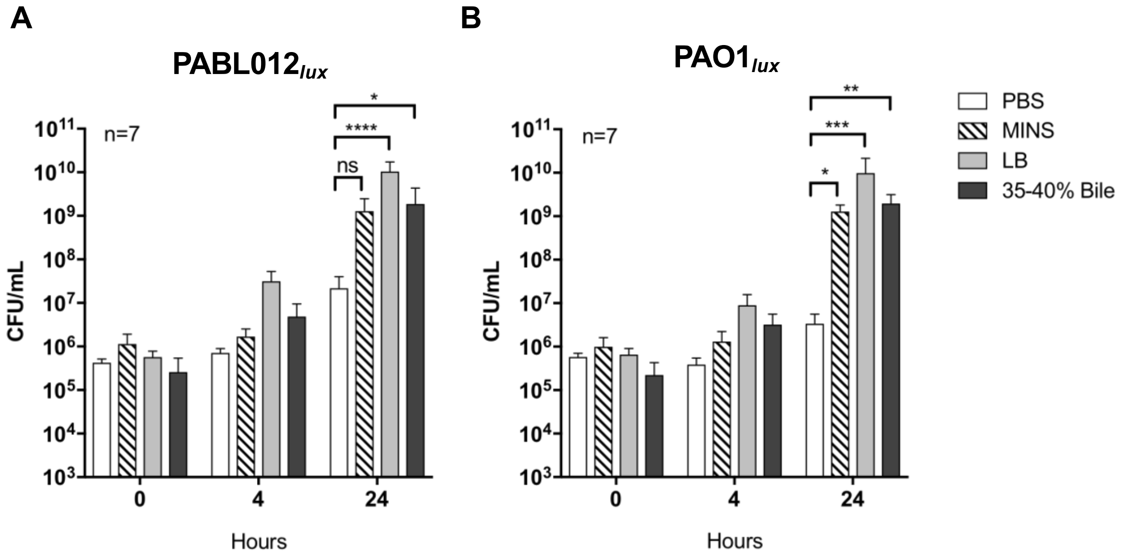


**Extended Data Figure 1. Bile promotes the replication of *P. aeruginosa*.** *In vitro* bacterial growth was enumerated for PABL012$_{lux}$ (A) and PAO1 (B) in PBS, MINS, LB, or PBS supplemented with 35-40% mouse bile at 0, 4, and 24-hour post-inoculation. Columns represent the mean CFU/mL with standard deviation (SD, whiskers) (n=7). P-values were calculated by ANOVA analysis with Kruskal-Wallis test correction for multiple comparisons. (ns) not significant, (*) $p < 0.05$, (**) $p < 0.005$, (***) $p < 0.001$, (****) $p < 0.0001$.

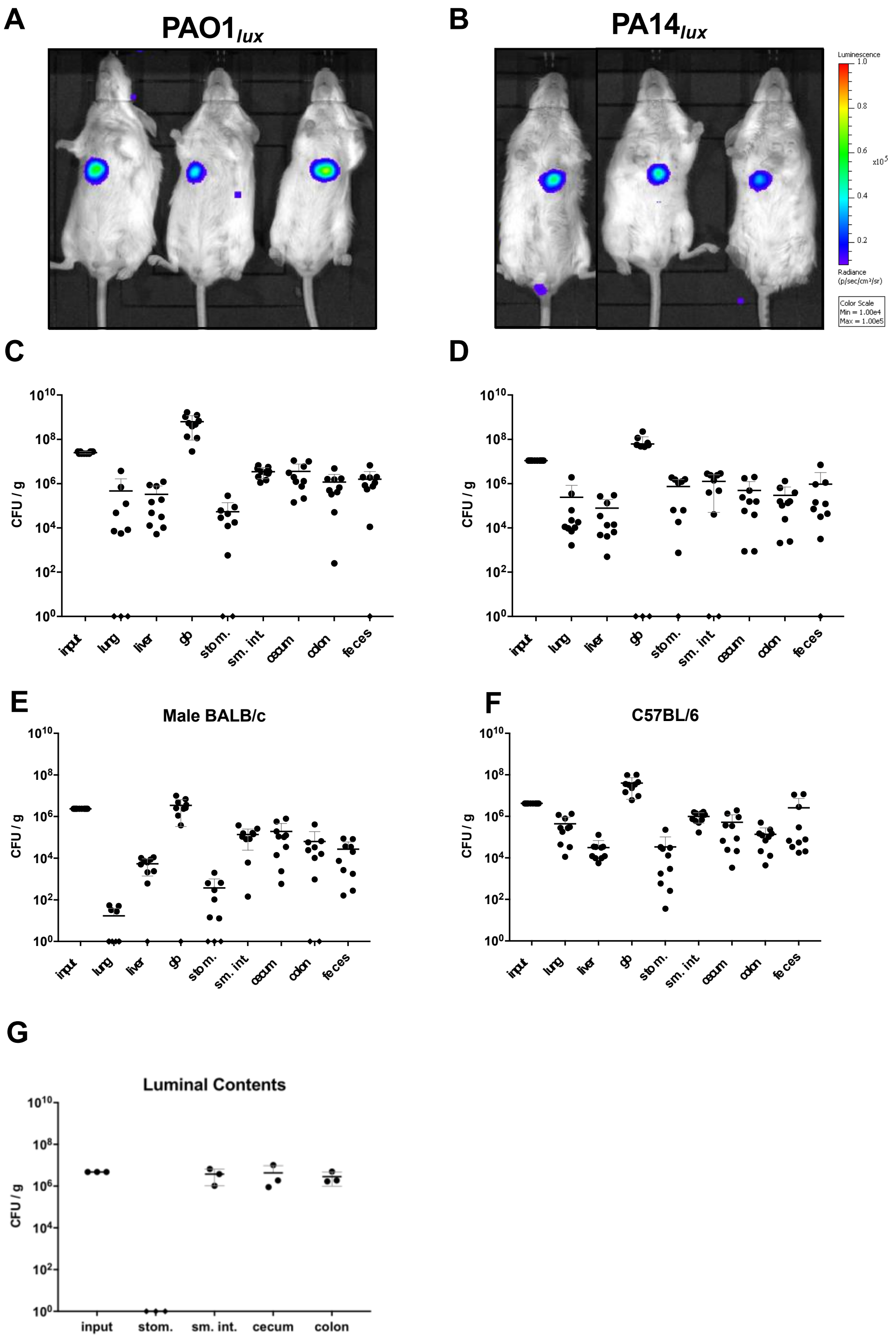


**Extended Data Figure 2. Gastrointestinal shedding of *P. aeruginosa* appears universal and independent of mouse genetics or gender.** BALB/c mice were infected intravenously with ~1 x $10^6$ CFU of PAO1 (A, n=5 x 2 replicates) and ~2 x $10^5$ CFU of PA14 (B, n=6 x 2 replicates) and imaged on the IVIS at 24 hpi. Mice were subsequently euthanized, and bacteria were enumerated from several anatomical sites. (C, D) Bacterial density as measured by CFU per gram organ [CFU/g] are presented (black circles). (E) Six- to eight-week-old male BALB/c mice (n=5 x 2 replicates) were infected intravenously with ~2 x $10^6$ CFU of PABL012*lux*, organs harvested as 24 hpi and plated for CFU enumeration. (F) Six- to eight-week-old C57BL/6 (n=5 x 2 replicates) mice were infected intravenously with ~2 x $10^6$ CFU of PABL012*lux*, organs harvested as 24 hpi and plated for CFU enumeration. (G) BALB/c mice (n=3) were infected intravenously with ~2 x $10^6$ CFU of PABL012*lux*, bacteria were enumerated from the luminal contents of the GI tract by extrusion of contents following dissection, and represented as CFU per gram. Each symbol represents one mouse; when no bacteria were recovered, mice are represented as diamonds on the x-axis. Geometric means (horizontal lines) and SD (whiskers) are shown. (gb) gallbladder, (stom.) stomach, (sm. int.) small intestine

**A**

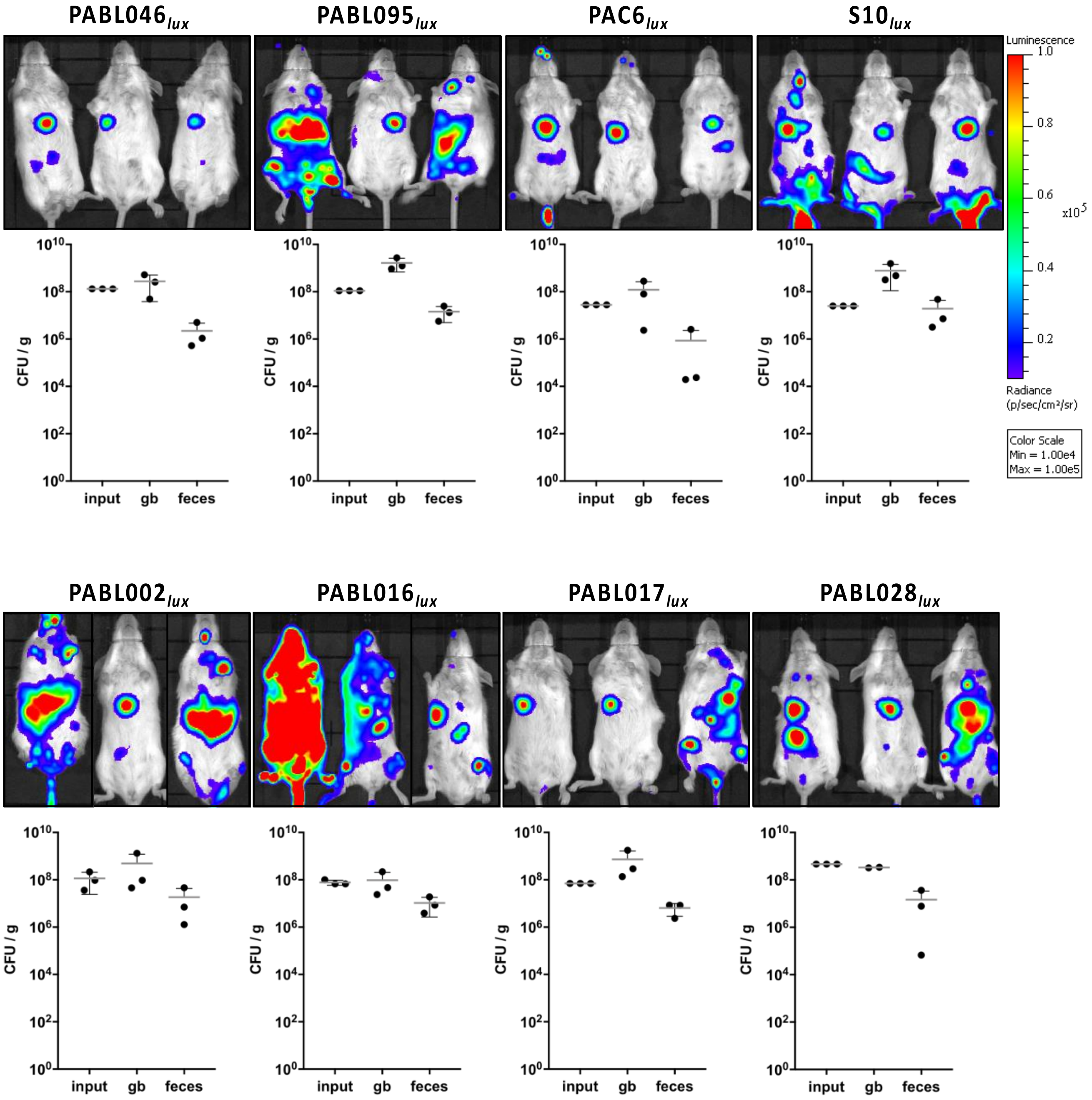
PABL046lux
PABL095lux
PAC6lux
S10lux
Luminescence
1.0
0.8
0.6
0.4
0.2
x10^5
Radiance
(p/sec/cm²/sr)
Color Scale
Min = 1.00e4
Max = 1.00e5
CFU / g
10^10
10^8
10^6
10^4
10^2
10^0
input
gb
feces
PABL002lux
PABL016lux
PABL017lux
PABL028lux

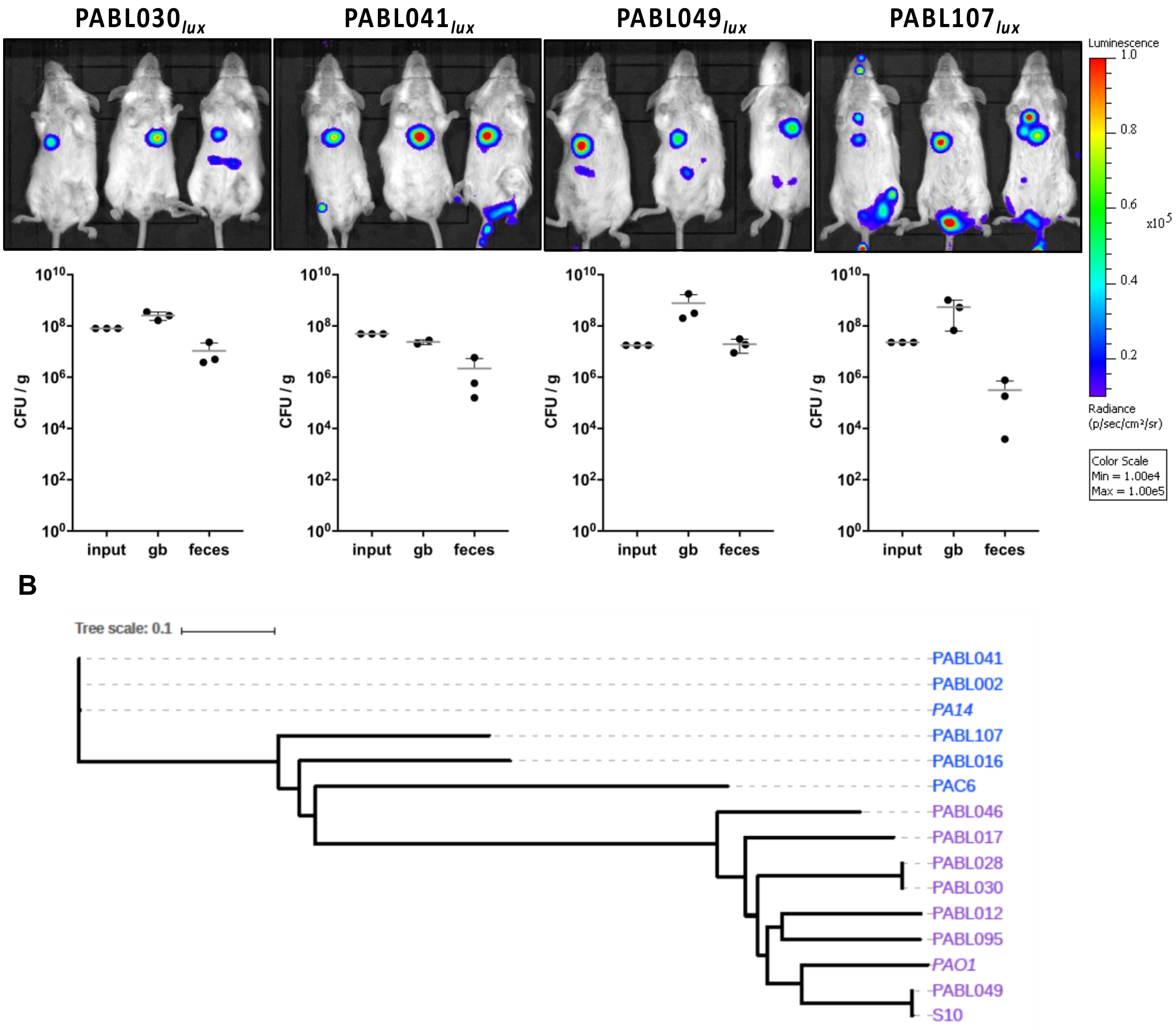


**Extended Data Figure 3. Dissemination to the gallbladder and fecal shedding are shared features among *exoS*- and *exoU*-containing clinical isolates of *P. aeruginosa*.** (A) BALB/c mice were infected intravenously with 8.8 x $10^5$ – 4.8 x $10^7$ (dose specific to each strain) of *P. aeruginosa* strains PABL046$_{lux}$, PABL095$_{lux}$, PAC6$_{lux}$, S10$_{lux}$, PABL002$_{lux}$, PABL016$_{lux}$, PABL017$_{lux}$, PABL028$_{lux}$, PABL030$_{lux}$, PABL041$_{lux}$, PABL049$_{lux}$, and PABL107$_{lux}$ and imaged at 24 hpi using IVIS. Mice were subsequently euthanized, and bacteria were enumerated from gallbladder and fecal homogenates. Log CFU per gram organ (black circles) of recovered bacteria are presented. Each circle represents one mouse (n=3, minimum 2 replicates). Geometric means (horizontal lines) and SD (whiskers) are shown. (B) A parsimony tree based on SNP loci present in 95% of the genomes was generated using kSNP v3.0.21[70] and visualized in iTOL[71]. The midpoint rooted tree was used to visualize *exoU*+ strains (blue) and *exoS*+ strains (purple).

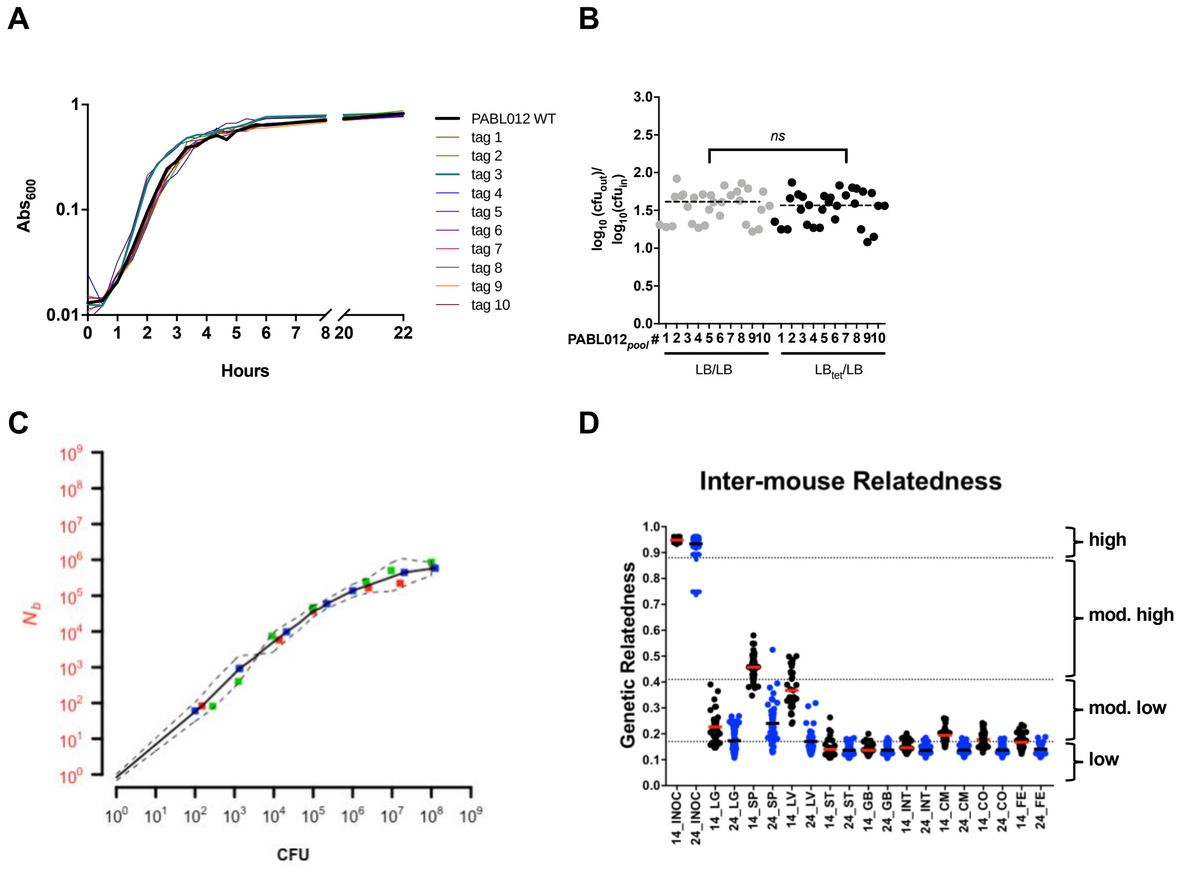


**Extended Data Figure 4**. **STAMP control experiments and calculations.** (A) *In vitro* growth of parental PABL012 and a selection of 10 individually barcoded PABL012*pool* (tag 1-10) strains in liquid culture over 22 hours. (B) The stability of barcode insertion was determined by comparing the output and input CFU for the same 10 individually barcoded strains grown initially in liquid culture without selection for the barcodes (LB) for 20 hours followed by growth on agar plates with tetracycline supplementation for growth of barcoded bacteria ($LB_{tet}$) or without tetracycline supplementation for growth of both tagged and untagged bacteria (LB). The data are presented as a ratio of output to input CFU (grey circle [LB vs. LB], black circle [$LB_{tet}$ vs. LB]). The dashed line represents the overall median for each condition where PABL012*pool* strains with tags #1-10 were tested in triplicate (each data point represents one replicate). No significant difference in the ratio of output to input with or without tag selection was detected ($p$ = 0.62, Mann-Whitney test). (C) Calibration curve for correspondence of mathematically derived founding population sizes ($N_b$; y-axis) with experimentally determined founding population sizes (CFU; x-axis) for three independent technical replicates (filled red, blue, green squares). The solid line indicates the median and the dashed black lines represent the 95% confidence intervals (CI). (D)

Genetic relatedness (GR) of the inoculums and bacteria from the same organ type in different mice are presented. For the inoculums, each circle represents the allelic frequency of barcodes in each inoculum compared to the allelic frequencies of the same barcodes in a different inoculum (technical replicates, n=30). For inter-mouse GR, each circle represents one organ in pairwise comparison with the same organ from a different mouse (14 hpi black, 24 hpi blue). Geometric means (14 hpi [red] and 24 hpi [black] horizontal bars) are shown. Dashed lines denote GR categories which are defined as high (≥0.88), moderately high (<0.88, ≥0.41), moderately low (<0.41, ≥0.17), and low (<0.17). LG (lung), SP (spleen), LV (liver), ST (stomach), GB (gallbladder), IN (small intestine), CM (cecum), CO (colon), FE (feces).

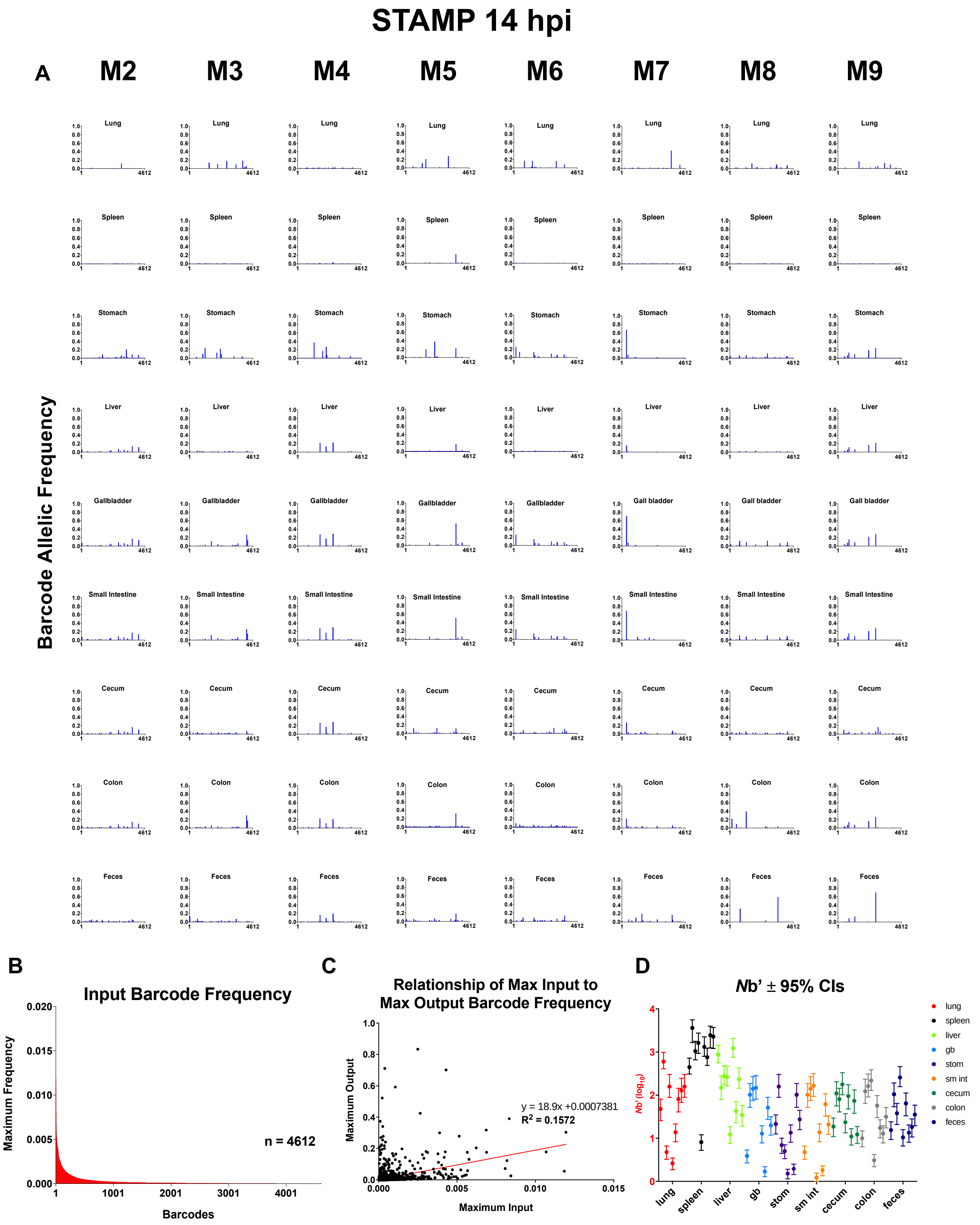


**Extended Data Figure 5. Analysis of individual STAMP barcode distributions across all organs in all mice at 14 hpi.** BALB/c mice (14 hpi, Mouse 2-9) were infected with *P. aeruginosa* strain PABL012*lux* by intravenous inoculation of ~2 x $10^6$ CFU and organs harvested as described previously. (A) Allelic frequencies of all unique barcodes are presented (y-axis) for every organ and every mouse (M2-9, M1 is shown in main text). All 4612 unique barcodes are arranged in identical order along the x-axis. (B) For all 4612 barcodes processed, the maximum frequency found in any inoculum (across all 30 technical replicates) is presented along the y-axis. Barcodes are arranged from most frequent (0.0119, 1.19%) to least frequent (0.000007, 0.0007%). (C) To determine the relationship between input frequency and output frequency for all 4612 barcodes, we plotted the maximum output frequency of that barcode (y-axis) in any mouse versus the maximum input frequency of that barcode found in the inoculum (x-axis). A linear regression was performed, revealing a poor correlation between the input barcode and the output barcode frequency ($R_2$ = 0.1572). (D) The calculated $N_b'$ values and their associated 95% confidence intervals (CI, whiskers) for every organ in every mouse at 14 hpi; circles, lung (red), spleen (black), liver (light green), gallbladder (light blue), stomach (purple), small intestine (orange), cecum (green), colon (grey), feces (navy).

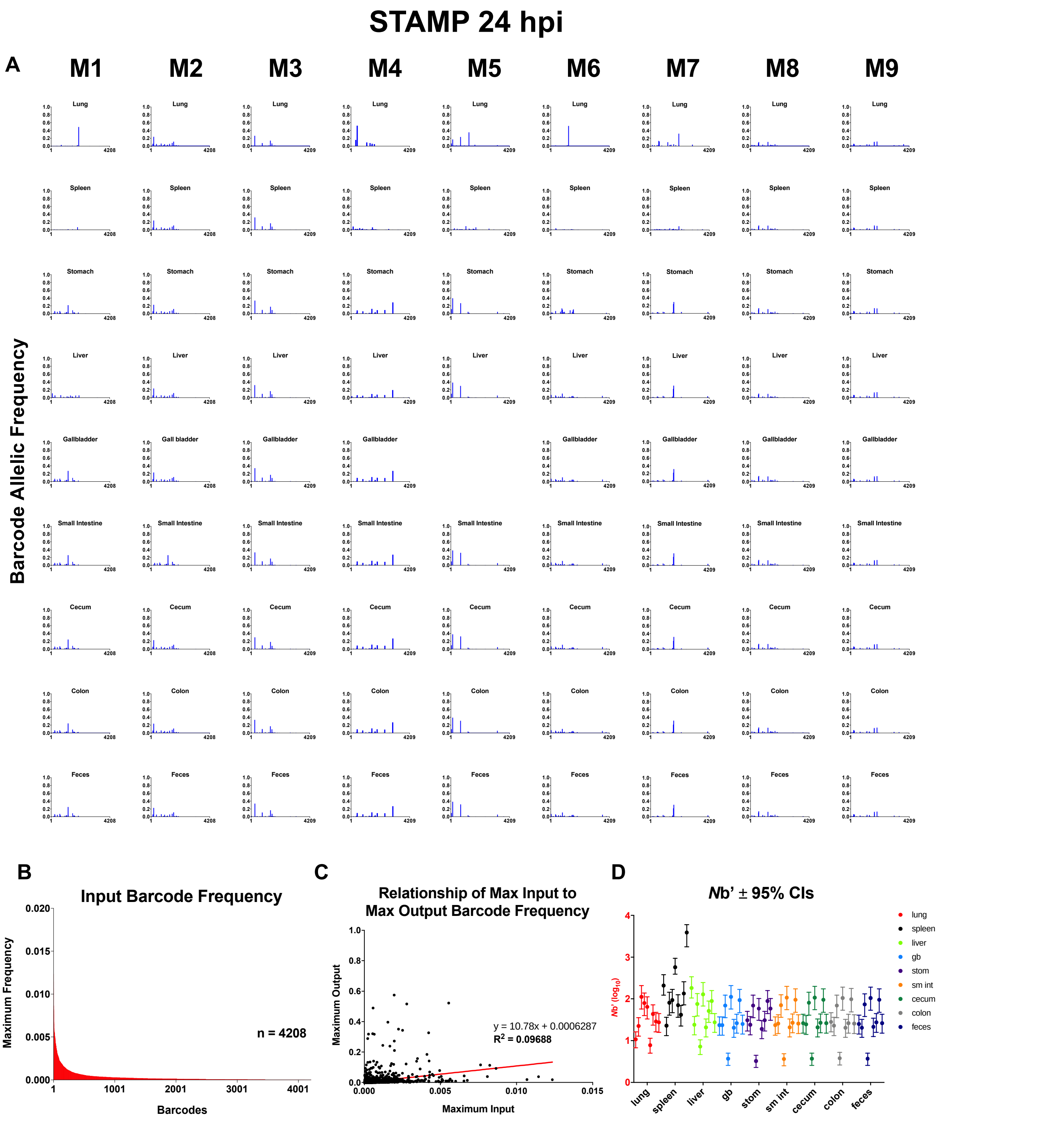


**Extended Data Figure 6. Analysis of individual STAMP barcode distributions across all organs in all mice at 24 hpi.** BALB/c mice (24 hpi, Mouse 1-9) were infected with *P. aeruginosa* strain PABL012$_{lux}$ by intravenous inoculation of ~2 x 10$_{6}$ CFU and organs harvested as described previously. (A) Allelic frequencies of all unique barcodes are presented (y-axis) for every organ and every mouse (M1-9, M10 shown in main text). All 4208 barcodes are arranged in identical order along the x-axis. Mouse 5's gallbladder was unable to be processed following dissection. (B) For all 4208 barcodes processed, the maximum frequency found in any inoculum (across all 30 technical replicates) is presented along the y-axis. Barcodes are arranged from most-frequent (0.0124, 1.24%) to least frequent (0.000011, 0.00011%). (C) In order to determine the relationship between input frequency and output frequency for all 4208 barcodes, we plotted the maximum output frequency of that barcode (y-axis) in any mouse versus the maximum input frequency of that barcode found in the inoculum (x-axis). A linear regression was performed, revealing a poor correlation between the input barcode and the output barcode frequency ($R_2$ = 0.09688). (D) The calculated $N_b'$ values and their associated 95% confidence intervals (CI, whiskers) for every organ in every mouse at 14 hpi are presented; circles, lung (red), spleen (black), liver (light green), gallbladder (light blue), stomach (purple), small intestine (orange), cecum (green), colon (grey), feces (navy).
